# Growth effects and the underlying genetic architecture of inbreeding depression in a wild raptor

**DOI:** 10.1101/2025.08.17.670740

**Authors:** Anna M. Hewett, Eléonore Lavanchy, Alexandros Topaloudis, Tristan Cumer, Anne-Lyse Ducrest, Céline Simon, Bettina Almasi, Alexandre Roulin, Jérôme Goudet

**Affiliations:** Department of Ecology and Evolution, University of Lausanne (UNIL), Lausanne, Switzerland; Swiss Institute of Bioinformatics, Lausanne 1015, Switzerland; Swiss Ornithological Institute, Sempach, Switzerland

## Abstract

Despite its potentially devastating effects, the prevalence and underlying mechanisms of inbreeding depression in wild populations are still relatively under-explored. Here, we use whole-genome sequence data from >3,000 wild barn owls from Switzerland to investigate the presence, severity, and genetic architecture of inbreeding depression in three morphological traits. Using a combination of linear models (accounting for age) and non-linear models (to measure growth effects) we clearly show inbreeding depression is present in this population. Moreover, by breaking-down the timing of the effects we also have a better ability to detect inbreeding depression, as in some traits we find that it manifests during juvenile growth, and in others during adulthood. To our knowledge this is the first study to show direct evidence for inbreeding depression during the crucial early life weight gain period in a wild animal. We further show that certain trait-specific patterns may reflect differences in environmental influences across life stages, as we find that heritability is often lower before adulthood. We also use two classes of genomic inbreeding coefficients: *F*_ROH_ and *F*_UniW_, and while the directionality of effects is equivalent, the strength of evidence regarding the presence of inbreeding depression differs depending on the coefficient used. This discrepancy might give some insight into the frequency distribution of responsible variants, as each coefficient weights variants differently based on their population frequencies. Finally, an assessment of local genomic inbreeding effects highlights a handful of regions with significantly deleterious effects, alongside many regions with a smaller contribution to the observed inbreeding depression. Overall, we provide a comprehensive overview of the effects of inbreeding in this wild population, highlighting the dynamic interplay between environmental influences and the selection pressure against inbred individuals.

## 2 Introduction

The study of inbreeding depression (i.e. the decreased fitness of inbred individuals) is crucial in fields from human genetics to conservation biology. In humans, inbreeding has been linked to a number of diseases, including Alzheimer’s disease (Vardarajan et al., 2015; Nalls et al., 2009; Ghani et al., 2015) and Schizophrenia (M. C. Keller et al., 2012). In endangered populations, where inbreeding can be intense and sometimes unavoidable, inbreeding depression limits population recovery and increases the risk of extinction (Kardos et al., 2023; L. F. Keller & Waller, 2002). However, despite its severity, the prevalence of inbreeding depression and its underlying mechanisms are still poorly understood in wild populations.

Detecting inbreeding depression in wild populations presents unique challenges. Such studies may require a large sample size to accurately detect inbreeding depression (M. C. Keller et al., 2012; Caballero et al., 2021; Lavanchy, Weir, & Goudet, 2024), unless the strength of inbreeding depression is exceptionally high. Moreover, merely obtaining phenotypic data and individual inbreeding coefficients required for studies of this nature can be difficult in natural populations. Obtaining phenotype data can be time-consuming and labour-intensive, but once collected it can prove invaluable in such studies. A major advance in the field was the use of genomic inbreeding coefficients, rather than those estimated from a population pedigree - although the latter is still a valid option. Often, genetic data is easier to acquire than an accurate pedigree in a wild population, and will provide estimates of realised inbreeding rather than predicted inbreeding. Hence, numerous genomic estimators of inbreeding coefficients have been established in recent years (reviewed in: Zhang et al. (2022)). These estimators are either based on the probability of identity-by-descent (IBD), and therefore range between 0 and 1 e.g. *F*_ROH_, or on the correlation between uniting parental gametes relative to the reference population (Wang, 2014), which can range between −1 and 1 e.g., *F*_GRM_, *F*_Uni_, and *F*_AS_. The choice of the most appropriate genomic inbreeding coefficient to study inbreeding depression has been widely debated (Yengo et al., 2017; Alemu et al., 2021; Caballero et al., 2021; Lavanchy, Weir, & Goudet, 2024), and is dependent on the demographic history of the focal population or the type of genotyping employed. Despite this, various genomic inbreeding coefficients have since been used to establish a reliable association between inbreeding and a number of fitness-related traits in various wild species, such as breeding success (Huisman et al., 2016), survival (Liberg et al., 2005; Kardos et al., 2023) and overall population growth (Bozzuto et al., 2019), highlighting their practical value in real-world conservation research. In addition, the acquisition of high-quality genomic data in recent years has renewed the interest in identifying loci responsible for observed inbreeding depression. For example, the contribution of (slightly) deleterious recessive loci versus over-dominant loci (Charlesworth & Willis, 2009), along with their effect size and distribution across the genome, still remains largely unknown in wild populations (Duntsch et al., 2023; Stoffel et al., 2021; Hewett et al., 2024).

Since early weight can be a key factor in determining juvenile survival, reduced growth rates are likely to be highly disadvantageous, particularly in a natural environment (Nielsen et al., 2012). The effect of inbreeding depression on growth rates has been relativity well explored in farmed species - where the daily growth of an individual represents significant economic gain (e.g. Gao et al. (2015); Nagy et al. (2013); Gholizadeh and Ghafouri-Kesbi (2016); Piles et al. (2023); Forneris et al. (2021); Silió et al. (2013)) - but few studies on growth rates in wild animals exist in the literature. One study on meerkats (*Suricata suricatta*) showed evidence for inbreeding depression in pup mass at emergence from the natal burrow, which is a good indicator of early life growth rates (Nielsen et al., 2012), although not a direct measurement of growth. For farmed species many studies measure individual growth at multiple fixed time points for uniformity, such as 8-week weight, meaning the growth rate can be easily calculated. However, in wild populations the exact age is often unknown prior to measurement, and repeated disturbances should ideally be kept to a minimum. Instead, studies on wild populations commonly obtain one morphological phenotypic measure per individual or, when multiple records exist, only one is taken during early juvenile growth stages. This highlights the need for further research to determine how widespread such effects may be.

The population of barn owls (*Tyto alba*) residing in south-west Switzerland have been the subject of a long-term study for over 30 years, and presents an ideal opportunity to investigate inbreeding depression in a wild population, especially during early growth rates. Since the last glacial maximum, barn owls have colonized Switzerland from glacial refugia in Portugal and Greece (Cumer et al., 2022). Now, they are wide-spread across Europe, but intense farming and homogenised landscapes has led to a population decline from which they may still be recovering (Frey et al., 2011). Throughout the breeding season, chicks and adults in installed nestboxes in the study area are routinely monitored (Béziers & Roulin, 2016). Regular phenotypic measurements are recorded, and blood samples are taken, usually when an individual is ringed. Because barn owls from this population are in an open landscape and can breed outside the study area, the pedigree for this populations is quite shallow, and inbreeding coefficients estimated from pedigree relatedness will likely underestimate the true level of inbreeding. Therefore, to more accurately capture individual inbreeding levels we use *>* 1 million variants from whole genome sequencing of *>*3,000 individuals to calculate genomic inbreeding coefficients (*F*_ROH_ and *F*_UniW_) and quantify inbreeding depression in three recorded morphological traits: bill length, weight and tarsus length. We also take advantage of these regular measurements covering a range of ages throughout the juvenile growth period, to investigate inbreeding depression in growth rates and identify the stages where these effects may be most pronounced. To further build on this, we disentangle the genetic architecture of inbreeding depression by associating local homozygosity-by-descent with these traits. Finally, we explore other recorded sources of variation in these traits and how these change over the life of an individual.

## 3 Methods

### 3.1 Study population, trait measurements and genetic sampling

Individual barn owls (*Tyto alba*) born within an area of south west Switzerland have been monitored as part of a long-term study since ~1986 (Béziers & Roulin, 2016; Frey et al., 2011; Roulin, Richner, & Ducrest, 1998). Currently there are 400 nest boxes at 316 sites (Supplementary Fig.1), which are visited regularly during the breeding season between March and October. When a clutch is found, the developmental stage of each egg is estimated. The first brood visit occurs when the oldest chick is predicted to be 30-days-old, based on a 32-day egg incubation time. Nestlings are then ringed with an individual identifier, have a blood sample taken, and are measured, including, but not limited to: bill length (millimetres), tarsus length (millimetres) and mass (grams), which are the three focal traits of this study. The age of each nestling is estimated based on their wing length at this first visit (Roulin, 2004). Between 2015 and 2024 instead of a visit at 30 days, two chick measurements were scheduled at 25 and 35-days-old. Final measurements are taken before fledging (~55-60 days) when the oldest chick is 55 days. Measurements are also taken of the parents, and when a parent has not been ringed previously (i.e. born outside of the study area) they are ringed, measured and have a blood sample taken.

As part of another experiment, in some years partial and full cross fostering of individuals between nests occurred (Roulin et al., 1998). In order to account for this in subsequent analysis we created a new variable which combined the ID of the born clutch and the raised clutch. For example, individuals born and raised in the same hypothetical clutch would have the clutch ID: 111 111, and those that were cross fostered would have a clutch ID: 111 222

A total of 3,085 individuals in the study population have currently been whole genome sequenced on a mix of high and low coverage. 502 individuals from the range of the species across the Western Palearctic (including 346 individuals from the focal population) were sequenced at high-coverage and used to build a reference panel, the remaining 2,768 individuals were sequenced at lower coverage and their full genotypes were imputed using the reference panel (see Topaloudis et al. and Cumer et al. for full details). To avoid any possible bias from sequencing depth we used only the 2,744 individuals sequenced on lower coverage (*<*5X) for our analysis. For estimation of genomic inbreeding coefficients, we used 37 high-quality autosomal super scaffolds with the synteny to the genome of the chicken *Gallus gallus* inferred in (Topaloudis et al., 2025). After phasing and quality control 10,074,749 high-quality called variants remained.

### 3.2 Estimating inbreeding coefficients

We estimated the inbreeding coefficient, *F*_UniW_, initially described in (Yang, Lee, Goddard, & Visscher, 2011) and modified in (Zhang et al., 2022) which measures the correlation between uniting gametes. *F*_UniW_ was calculated using the *get.funiw()* function in the JGTeach package (https://github.com/jgx65/JGTeach) as:

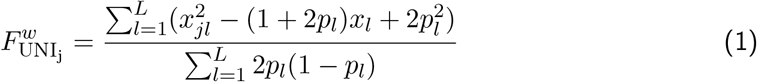

Where, *x*_*jl*_ ∈ {0, 1, 2} is the minor allele count for individual *j* at locus *l* and *p*_*l*_ is the allele frequency of the minor allele at the same locus.

We also estimated the runs of homozygosity (ROH) based inbreeding coefficient, *F*_ROH_. Runs of homozygosity of a given length were inferred through homozygous-by-descent (HBD) segments called using the RZooRoH package (v.0.3.1) using the genetic positions of SNPs (i.e. taking into account recombination rate) (Druet & Gautier, 2017, 2022) with 13 HBD classes and 1 non-HBD class with rates (R) of 2, 4, 8, 16, 32, 64, 128, 256, 502, 1024, 2048, 4096 and 8192 for the HBD classes and 8192 for the non-HBD class (Druet & Gautier, 2017). We then averaged the probability of belonging to a HBD segment in order to estimate inbreeding coefficients with the *cumhbd()* function, using a T value of 1024 (meaning that we only consider HBD segments coalescing less than 512 generations ago as identical-by-descent).

### 3.3 Statistical models

We ran all statistical models in brms v2.21.0 (Bürkner, 2017, 2018, 2021) in R v4.3.2. Trace plots, Rhats, and effective sample size measures were inspected for proper chain convergence. See Supplementary Tables 1-4 for specific details on warm-up iterations, total number of iterations, thinning intervals and specific sample and record numbers for each model. All model code, including priors used, is available at https://github.com/annamayh/Owls/tree/418fe0c27a0114795f8e6d175d917a617581a663/Inbreedingdepression.

#### Inbreeding depression in traits

To estimate the effects of inbreeding depression on three morphological traits we ran a standard repeated measures linear mixed model (LMM) with either bill length, body mass or tarsus length as the response variable. For all three traits we fitted age of the individual as a fixed effect modelled as a Gompertz growth curve,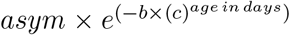S, where *asym* is the asymptote, *b* is the initiation of the growth curve and *c* is the growth slope. A new variable was created using the population average estimates of *asym, b* and *c* and the individuals age in days at the time of measurement (See Supplementary Figs.2, 3 and 4 for parameter estimates and curve). Sex and rank of the individual (i.e. numeric order of individuals born within a clutch) were also fitted as a fixed effects. Additional random effects were: observer, birth year of the individual (as factor), birth month of the individual (as numeric), individual nestbox location, individual clutch identity, and finally, the individual permanent environment to account for repeated measurements. We also included a beta dosage genomic relatedness matrix (GRM) as a random effect, in order to account for population and family structure as demonstrated in Lavanchy, Weir, and Goudet (2024). Finally, we fitted a genomic inbreeding coefficient, either *F*_UniW_ or *F*_ROH_ as defined above, as a fixed effect to estimate the effect of inbreeding coefficients on the trait value (i.e. inbreeding depression)

#### Inbreeding depression in growth rates

Given the availability of multiple records for each individual over its lifespan, particularly during the juvenile growth period, we then further explored at what point in an individuals life inbreeding depression may be most severe. To determine the effect of inbreeding on growth we estimated the effect of inbreeding coefficients on a three parameter version of the Gompertz growth curve (commonly used to model biological and economic growth (Tjørve & Tjørve, 2017)), using a hierarchical non-linear model with the following structure:

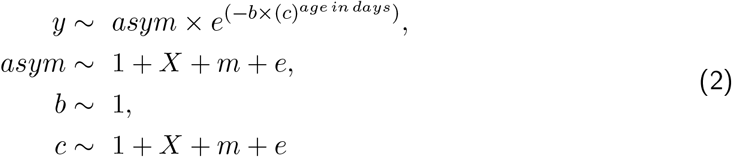

Where, in this case *age in days* is the age of the individual, *asym* represents the asymptote of the trait, *b* the initiation of the growth curve and *c* the growth slope. Here, *X, m* and *e* represent the fixed, random and residual effects on each parameter respectively. We used this particular re-parametrisation of the Gompertz growth curve with one exponential. During refinement of the model we also tested the double exponential Gompertz growth curve and a logistic growth curve. The former failed to converge and the latter was not a good representation of the data. For all three traits we fixed the intercept of *b*, so assuming that no fixed effect variable had an effect on the initiation of the growth. This was to aid model mixing and, as the age of an individual is an estimate (see Methods 1), effects on this specific parameter may be confounded whereas other parameters will be less affected. In each trait, certain modifications were made to optimise convergence times: for bill length we exponentiated the *c* parameter to prevent estimates from being below zero (impossible for a growth parameter), and for tarsus length, two asymptotes were used to allow the growth curve to start further from the origin i.e.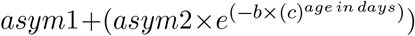. *X* (fixed effects) and *m* (random effects) for the *asym* parameter were the same as in the LMM described above, excluding the GRM to aid convergence. During model refinement co-variates affecting the growth rate parameter (*c*) were simplified to allow for model convergence, whereby, variables were removed if they could feasibly have little impact on individual growth and showed to have minimal effect on the results. This led to inbreeding coefficient, rank and sex of the individual as fixed effects and individual ring identity as a random effect to account for repeated measurements.

#### Heritability in stages

To further disentangle other sources of genetic and environmental variation over different life stages, we firstly split records into juvenile stages (≤180 days old) and adult stages (*>*180 days old) as most barn owls have fledged by ~ 90 days and can breed the following year *>* 6 months later (Béziers & Roulin, 2021). Then, we ran a repeated measures animal model with the following model structure:

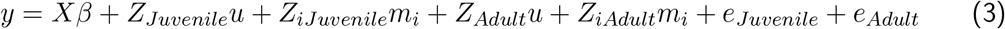

Where, *y* is a vector of the trait measurements and *X* is an incidence matrix relating individual measures to the vector of fixed effects, *β. Z_Juvenile_* and *Z*_*iJuvenile*_ are incidence matrices relating individual juvenile measures to additive genetic and the other remaining random effects respectively. In contrast, *Z*_*Adult*_ and *Z*_*iAdult*_ are incidence matrices relating individual adult measures to additive genetic and other remaining random effects. *u* is the relatedness matrix estimated from genomic data (GRM) as above, and *m*_*i*_ is a vector of additional random effects. Finally, *e*_*Juvenile*_ and *e*_*Adult*_ are vectors of residual effects in the juvenile and adult stages respectively. We used the same fixed and random effects described in the LMM, with the exception of rank being treated as a random effect here.

We calculated narrow sense heritability per stage as:

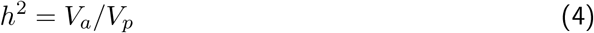

Where, *V*_*a*_ and *V*_*p*_ are the additive genetic variance and phenotypic variance respectively, for each growth stage (juvenile or adult). We removed the variance created by the observer in our estimates of *V*_*p*_ as these are biases created through sampling with little biological interpretation and may lead to an underestimation of narrow sense heritability (Hoffmann, 2000), although they are still necessary to account for within the model.

#### Genetic architecture of inbreeding depression

We firstly calculated local HBD probabilities using the *probhbd* function in RZooRoH for non-overlapping windows of 2500 SNPs (~ 100 kbp assuming 1 SNP every 40 bp on average). Then, for each window we ran separate LMM including both the local HBD probability and the genome-wide *F*_HBD_ as fixed effects. We repeated this process using local *F*_UniW_ calculated from a modified version of the *get.funiw()* function from the JGTeach package (https://github.com/jgx65/JGTeach) to estimate *F*_UniW_ per window rather than genome-wide. All other fixed and random effects were kept the same as the previous models with the exception of the GRM which was omitted to aid model convergence as during refinement it was shown to have a minimal effect on the estimates. P-values for windows were calculated as the proportion of post-warm up draws that were of the opposite value to the mean estimate of the effect of inbreeding. To account for the 4,010 independent tests for each window we used a Bonferroni corrected threshold of 1.25 *×* 10^−5^, where any window with a p-value below this threshold was then treated as a significant window.

## 4 Results

### 4.1 Evidence for inbreeding depression in morphological traits

We first estimated inbreeding depression in bill length, tarsus length and body mass with two different inbreeding coefficients, either: weighted *F*_UniW_ (Zhang et al., 2022) or *F*_ROH_ (McQuillan et al., 2008; Druet & Gautier, 2017), using a standard linear mixed model (LMM). Next, we further explored when inbreeding depression may be most severe. Rather than simply accounting for the age of an individual (as above), we ran a hierarchical non-linear mixed model (NLMM) to investigate the effect of inbreeding on growth.

For bill length, we showed consistent evidence for inbreeding depression in the LMM, regardless of whether *F*_ROH_ or *F*_UniW_ was used as the inbreeding coefficient Fig.1B. An individual with an inbreeding coefficient of 0.25 (i.e. the product of a father-daughter or full-sibling mating) is predicted to have a bill length ~ 3.1% and 5.2% shorter for *F*_UniW_ and *F*_ROH_ respectively in comparison to an outbred individual (F=0), Supplementary Table 5 and 6. The NLMM with *F*_UniW_ further suggests that these differences are most pronounced after growth. That is, inbred and outbred individuals do not differ considerably during the hatchling/nestling period (0 - ~ 30 days) but their ‘final’ bill length is different. Moreover, we observe that the estimate is within the same magnitude as the estimate from the LMM - ~2.4% shorter, Supplementary Table 17. However, counter to expectations, this was not the case using *F*_ROH_ in the NLMM where no parameters showed a significant effect of *F*_ROH_, despite finding evidence with the LMM Fig. 1F and Supplementary Table 18.

**Figure 1:**
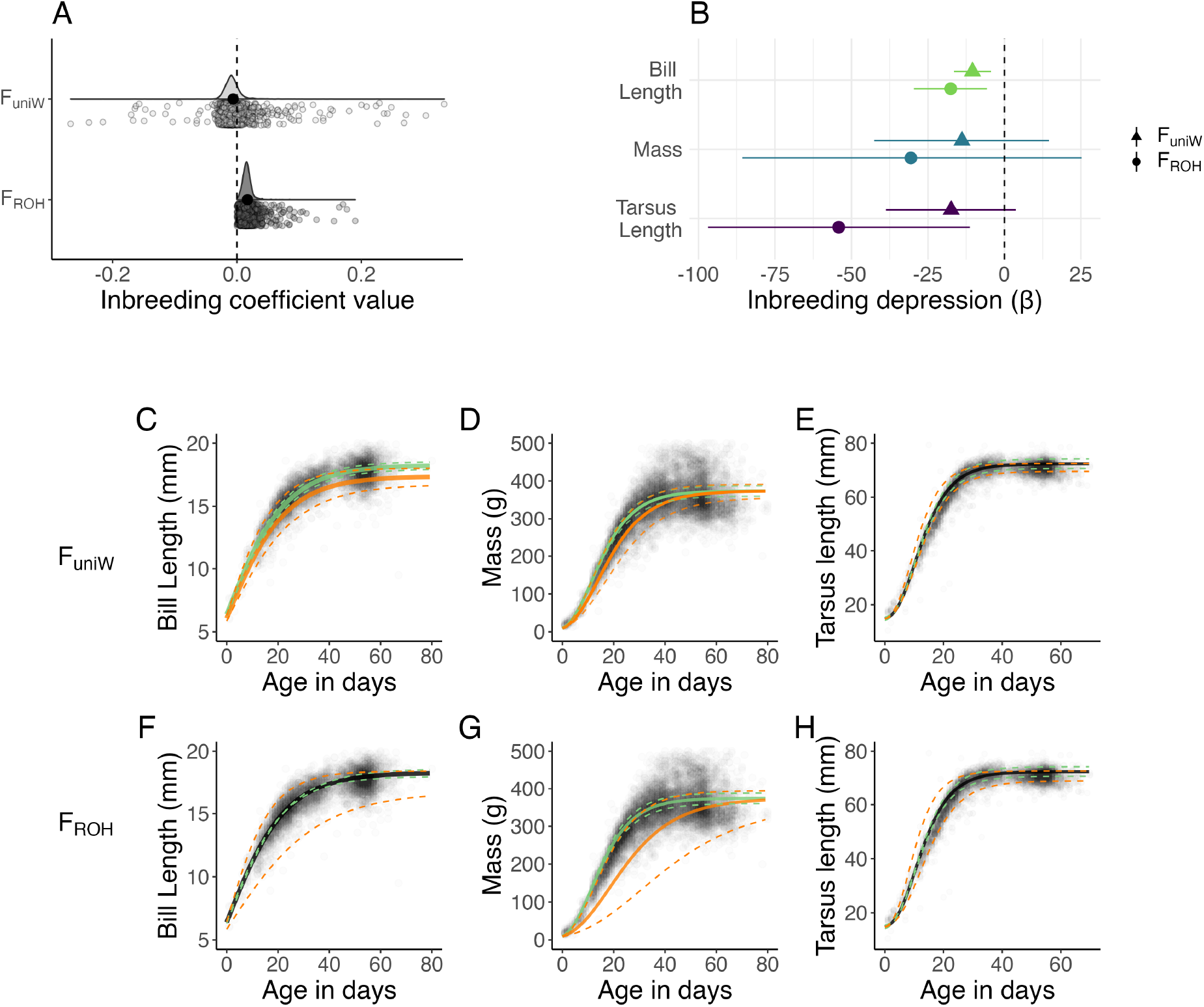
**A**: Distribution of two inbreeding coefficients **B**: Inbreeding depression (*β*) estimated from linear mixed models (LMM) in three morphological traits of barn owls using two inbreeding coefficients: *F*_UniW_ or *F*_ROH_. Mean posterior estimate is shown by the symbols and 95% credible intervals are indicated by the corresponding lines around the mean estimate, used to determine the significance. Estimates are relative to the scale of the trait and show the difference between an inbred (F=0.5) and and an outbred individual (F=0). Note that for visualisation purposes all traits are plotted using the same x-axis but the scales differ between traits e.g. the mean bill length is 16.5mm (scaled to 3 digits), the mean mass is 323g and the mean tarsus length is 65.4mm (scaled to 3 digits). **C-H**: Inbreeding depression in growth rates of the three traits and two inbreeding coefficients estimated from non-linear mixed models (NLMM). The three models where the 95% credible intervals for the estimate of inbreeding coefficient did not overlap zero are coloured (C, D and G), with the predicted growth curve for inbred individuals (F=0.5) shown in red and an average individual (F=0) shown in green. Predicted slopes are overlaid on top of raw data points. The x-axis is truncated at an age when the growth asymptotes but estimates and data extend beyond the axes.

For mass, using the LMM we found no statistical evidence for inbreeding depression, Fig.1B. While the estimates of inbreeding coefficients showed an overall negative trend with the mass of an individual, suggesting inbred individual may be lighter, the 95% credible intervals overlapped zero for both *F*_UniW_ and *F*_ROH_, Fig.1B, Supplementary Tables 7 and 8. In contrast, the NLMM showed a significant effect of individual inbreeding coefficients (*F*_UniW_ and *F*_ROH_) in the early juvenile weight gain period (growth parameter *c*), Fig. 1D & G and Supplementary Table 20. For example, an inbred individual with an *F*_UniW_ of 0.25 was predicted to grow ~6% slower in the first 25 days of life, gaining on average 273g compared to 290g in its outbred counterpart. Using *F*_ROH_, we predict a more pronounced difference, with inbred individuals growing ~17% slower, gaining 250g compared to 293g over the same period.

Finally, for tarsus length, we found inconsistent results between the inbreeding coefficient used in the LMM. Both *F*_UniW_ and *F*_ROH_ in general indicated that higher inbreeding coefficients lead to a smaller tarsus length, although only *F*_ROH_ showed a significant association, Fig 1 and Supplementary Tables 9 and 10. An individual with *F*_ROH_ equal to 0.25 is estimated to have a tarsus length ~4.5% smaller than a completely outbred individual. However, using the NLMM we found no association with *F*_ROH_ or *F*_UniW_ for growth rate (*c*) or the asymptote, Fig. 1 E & H and Supplementary Tables 21 and 22.

Regarding the other fixed effects in the model, using the LMM we find that females are heavier, have longer bills but a shorter tarsus, Supplementary Tables 5 - 10. We also find that lower ranked individuals (i.e. those born later than their siblings) are lighter and have shorter bills and tarsi, Supplementary Table 5 - 10. We found no association of rank or sex on any growth rates (*c*) and our results for the ‘final’ asymptotic weight reflect those in the LMM (Supplementary Tables 17 - 22), with the exception of lower rank individuals showing a heavier final weight - likely because only the heavier individuals at a lower rank survived.

### 4.2 Heritability changes over time

Understanding the sources of variation that contribute to each trait over the course of an individual’s life may give more context to the above analysis. Estimates of heritability increased between juvenile and adult stages for all traits, although this increase was only significant in mass, Fig.2 and Supplementary Fig.5. The proportion of variance explained by additive genetic variance (i.e. trait heritability, *h*^2^) in juveniles was 0.356, 0.06 and 0.157 for bill length, mass and tarsus length respectively. In contrast, the equivalent *h*^2^ for adults were: 0.364, 0.419 and 0.216 (Supplementary Tables 23-25). Evolvability (additive genetic variance divided by the trait mean squared) also significantly increased between juvenile and adult mass, but decreased for the other two traits (Supplementary Fig.5).

**Figure 2:**
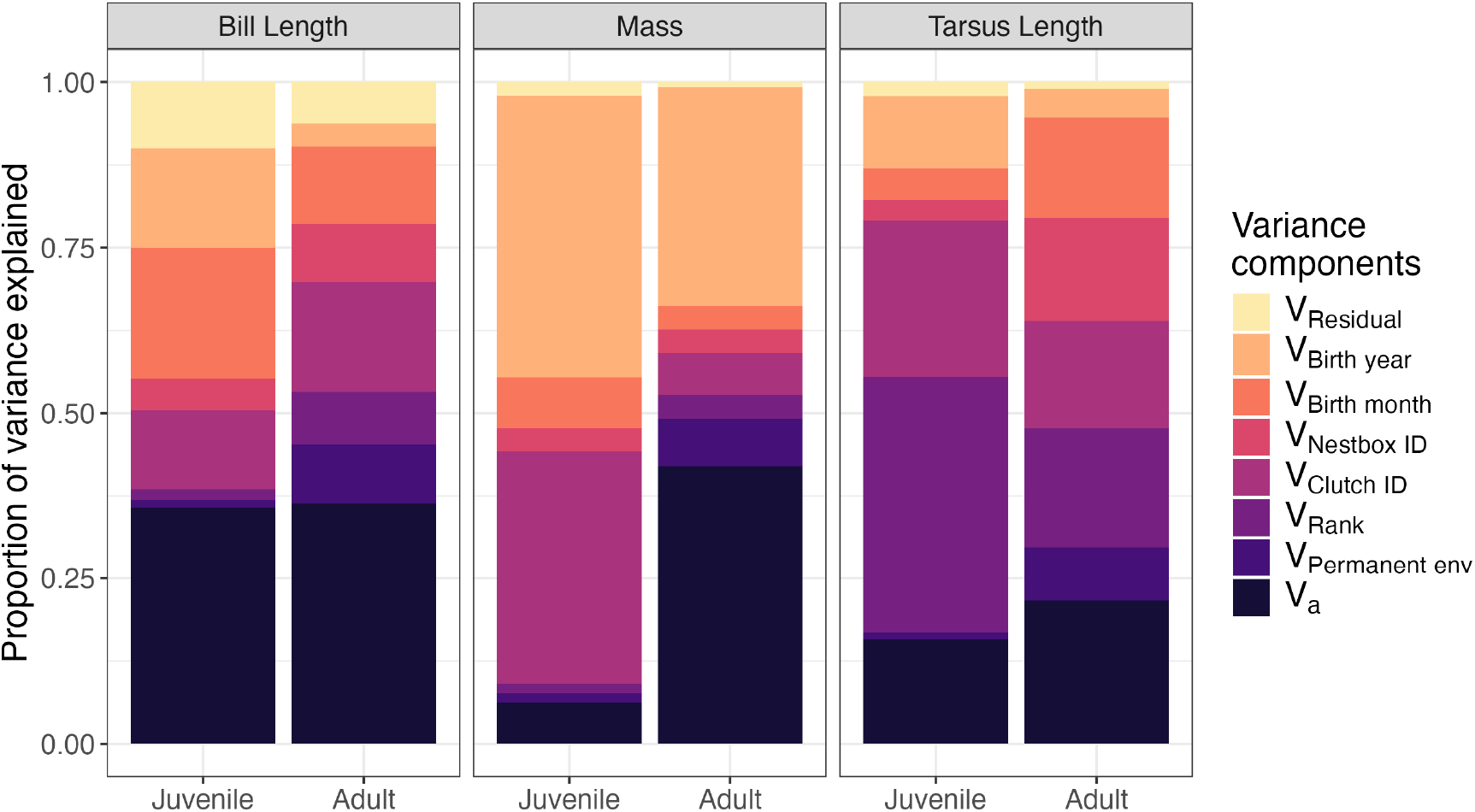
Proportion of variance explained by different components across the juvenile (≤180 days old) and adult stages (*>*180 days old) for three morphological traits. Thus, additive ge-xnetic variance (Va) in dark purple (bottom) gives an indication of the narrow sense heritability for each trait at different life stages.

In juveniles, environmental factors such as birth year and clutch identity consistently had a greater contribution to the overall phenotypic variance compared to their contribution to adult phenotypic variance (Fig.2). For mass in particular, the birth month and year as well as the clutch identity, explains the majority of the phenotypic variance in juveniles, likely due to fluctuations in food availability within and between years and the level of parental investment. In contrast, for adults, in all traits additive genetic variance was the largest contributor to overall phenotypic variance, exceeding the contribution of any other environmental variance components (Fig.2).

### 4.3 Genetic architecture of inbreeding depression

In order to uncover the genetic loci underlying inbreeding depression, we calculated local *F*_ROH_ across non-overlapping windows each containing 2500 SNPs (~ 100 kb assuming 1SNP/40bp).

We use *F*_ROH_ as a measure of local inbreeding depression in this analysis. RZooRoH provides a convenient framework to calculate these metrics, quantifying the probability of belonging to an HBD segment for any given window size. For each window we ran a separate LMM fitting both the local *F*_ROH_ and the genome-wide *F*_ROH_ to determine the localised effect of inbreeding depression. A similar approach was used in Stoffel et al. (2021) and Duntsch et al. (2023) although here we use a Bayesian framework. We also ran this analysis using local *F*_UniW_, yielding dissimilar results (Supplementary Fig.7). We continued with *F*_ROH_ because unlike other estimates - such as a binary classification of the presence of a ROH or possibly *F*_UniW_ - using a local model based estimate of *F*_ROH_ incorporates any fine-scale uncertainty or bias existing from genotyping errors or ascertainment bias (Bertrand et al., 2019).

After accounting for multiple testing, we find nine windows with significantly deleterious effects (negative estimates) and three windows with significantly beneficial effects (positive estimates) across our three phenotypic traits, Fig 3 and Supplementary Tables 27 - 29. In bill length - which has the strongest and most consistent estimate of inbreeding depression in previous analyses - three windows are significantly deleterious on chromosomes 8, 10 and 17; while one window is significantly beneficial on chromosome 2E. We also find a marginally negative skew of effects sizes with the average effect of local inbreeding being slightly negative (mean and median window estimates ∼0.12 and ∼0.08 respectively and see Fig 3 right). In contrast, for body mass, where we did not detect inbreeding depression using the LMM in the previous analysis, three significant windows were detected — two with beneficial effects on chromosomes 9 and 15, and one deleterious on chromosome 4A (Fig 3). Here the mean effect size is slightly positive (0.01) and the median is negative (−0.01) but the distribution mainly centres around zero with limited skew. Finally, for tarsus length, for which inbreeding depression was detected with only *F*_ROH_ in the previous analysis, we identified five windows with deleterious effects, occurring on chromosomes 2A, 2C, 4B, 7 and 23. Interestingly, the average effect size of windows are positive (mean and median window estimates 0.13 and

**Figure 3:**
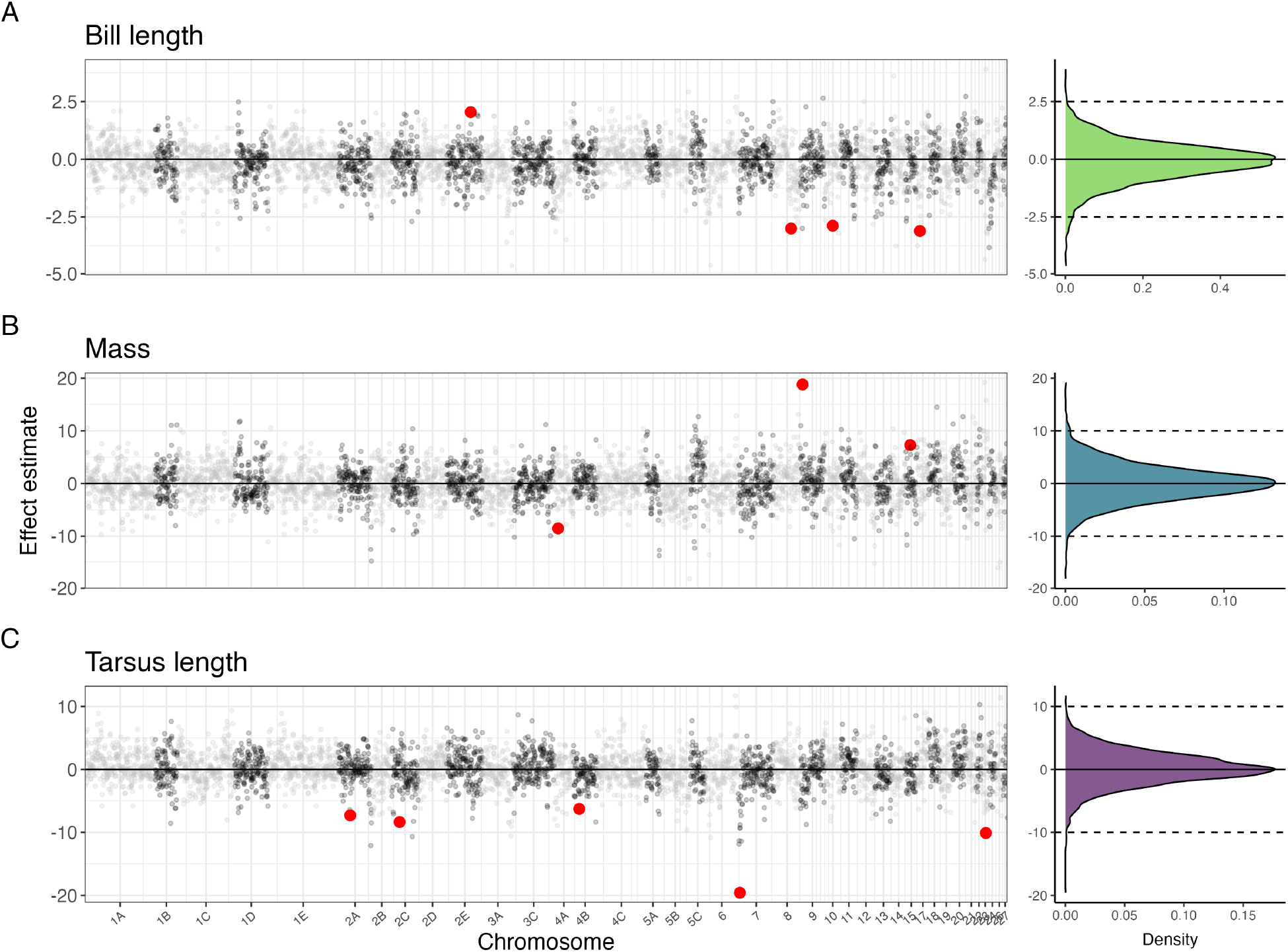
Genetic architecture of inbreeding depression. Left: Each dot represents the estimated effect of local inbreeding on the trait within a 2500 SNP window when local and genome-wide inbreeding *F*_ROH_ are fitted together in the same model. Chromosome numbers are based on the synteny between identified super scaffolds and the genome of chicken *(Gallus gallus)* in Topaloudis et al. (2025). Red coloured dots show significant windows crossing the Bonferroni corrected significance threshold where the p-value is calculated as the number of post-warm up iterations with an opposite values to the mean estimate (see methods). Solid line shows zero effect. Significant windows below the zero line indicate significantly deleterious windows - i.e. higher local inbreeding leads to a significantly reduced trait value. Significant windows above the zero line show significantly beneficial windows - where local inbreeding increases trait values Right: Density plots of estimated local inbreeding effects, again, solid line shows zero effect.

0.11 respectively), but the distribution is negatively skewed and leptokurtic, Fig 3 right. This indicates a number of outliers with a deleterious effects contributing to inbreeding depression in this trait. Although, the effect sizes of significant windows are still rather modest. An individual fully identical-by-descent(F=1) for the most highly associated window in tarsus length on chromosome 7 would have a tarsus ~2mm smaller. Given the average tarsus length of an adult is 70mm, labelling this regions as ‘severely deleterious’ may therefore be a slight overstatement of its impact. Nevertheless, the genetic architecture of inbreeding depression appears variable across our three traits and highlighting the importance of considering both widespread and localized effects of inbreeding.

## 5 Discussion

Using a long-term dataset of the Swiss barn owl population, we reveal clear evidence for inbreeding depression in three morphological traits: bill length, tarsus length and mass, deepening our understanding of inbreeding depression in the wild. While life-history traits such as survival and fecundity are generally expected to exhibit stronger signs of inbreeding depression (DeRose & Roff, 1999), they can be difficult to measure in a natural population, often requiring individuals to be monitored throughout their lives. In contrast, morphological traits are easily quantifiable and can be a good indication of inbreeding depression in wild populations (Bérénos et al., 2016; Huisman et al., 2016). Our findings align with previous work from just a handful of studies in wild birds showing that inbred individuals are lighter (Hajduk et al., 2018; Niskanen et al., 2020), have smaller skeletal sizes (Kruuk et al., 2002), and shorter bills (Niskanen et al., 2020). Such inbreeding depression can directly impact overall fitness as smaller individuals are less likely to survive in a natural setting. Therefore, the inbreeding depression we find in these three morphological traits, likely reflects a broader reduction in fitness.

We further uncover which stage within early development inbreeding depression is most severe combining linear (LMM) and non-linear (NLMM) mixed model approaches (Fig.1). We show that inbred individuals appear to be slower at gaining boody mass (Fig. 1 D and G). To our knowledge, this is only the second study to show evidence for inbreeding depression in early growth rates in a wild population. In the other, Nielsen et al. (2012) demonstrate inbreeding depression in pup mass at emergence, inferring the growth rate prior to emergence. Similar early life effects have been shown in domesticated birds (Rahmanian et al., 2015; MacLaury & Johnson, 1971), but it was previously unknown whether wild birds would also be affected at such an early life stage, particularly as the inbreeding level is often lower than in artificially selected populations. If these effects are widespread this presents a concern in conservation of small, endangered species, as early life weight and weight gain is expected to have a large impact on survival (Nielsen et al., 2012). Interestingly we also find that inbred individuals ultimately reach a lower asymptotic bill length but show no significant difference during the juvenile stage - although this is only true using *F*_UniW_ and not *F*_ROH_. Rather than any biological explanation, a more likely rationale is that measuring the length of a juvenile bill can be challenging due to its small size, reflected in the residual variance of bill length, Fig.2. Therefore, in this trait we likely have more power to detect inbreeding depression in adults, where the measurements are more accurate.

We should also highlight the role that selective disappearance may be playing in our results. For consistency, all adult records are from those that were also caught as juveniles, however, not all juveniles return to the study area once they fledge and it is unknown whether they survived or not. Thus, hypothetically, a small inbred individual measured during its growing stages but not surviving to the following year, will not have an adult measurement. Therefore, the majority of the adult data points in this analysis are for those that have already survived the first selection process. Moreover, because nestbox visits are centred around sampling clutches, the ratio of adults to juvenile measurements will be biased towards juvenile measurements. Hence, although the asymptote is partly determined using the juvenile records, a large proportion will be determined by the re-caught adults the following field season, of which there are also fewer, lowering our power to infer inbreeding depression, particularly in the asymptote.

Overall, our mixed approach using both LMMs (to account for age) and NLMMs (to break down stages of the growth curve) is the best technique to maximise the statistical power of our analyses. For bill length and body mass using the NLMM showed at which life stage the severity of inbreeding depression (or our ability to detect it) is the strongest. In tarsus length, we show inbreeding depression using the LMM (for *F*_ROH_ only), but this is not consistent in the NLMM. In this case, breaking down the trait into two parameters (asymptote and growth slope) reduced the power rather than heightening it. In future studies, where such records are available, we recommend using both methods to avoid missing key evidence for inbreeding depression.

Our results in Fig.2 may provide further context for understanding our findings in Fig.1. For all traits, we show that heritability is larger in adults than in juveniles - most clearly seen, and only significant, in body mass (Fig.2 and Supplementary Fig.5). This increased heritability with age is an established pattern, usually attributed to the decrease in non-genetic sources of variation, rather than increased additive variation (Wilson et al., 2007). In our study system, juvenile barn owls rely entirely on parental care for food and protection until fledging, making them more susceptible to environmental variation during early life, such as food availability in the year they are born (Fig.2). Therefore, competition for food during early life may be heightening inbreeding depression at this time - an inbreeding depression-by-environment interaction. A contrary, but not entirely unrelated explanation, again highlights the role of selective disappearance in our results. We also show a significant increase in evolvability (additive genetic variance over the squared trait mean) of mass (Supplementary Fig.5), supporting the idea that additive genetic variance is increased in adults. Hence, given there is a known selective advantage for larger body size (Kingsolver & Pfennig, 2004) and we show the possibility for heightened selection in adults, potentially many of the smaller inbred individuals do not survive to adulthood. In this case, we would not detect inbreeding depression in the adults, or in their asymptotic body mass.

Another related issue in this analysis may be a result of ascertainment bias. Our study utilises data from another project where a subset of individuals from the whole population were genotyped, prioritising individuals with extensive phenotype records. Given that we expect highly inbred individuals to be less likely to survive (and therefore have less records), individuals with more extreme inbreeding coefficients may be missing from our dataset. In addition, there may be some discrepancies in the predicted age of very young chicks. The age of an individual is predicted based on the wing length at the first visit (see methods). This is done to reduce the impact of visits on the growing chicks. However, given that we show inbred individuals tend to be smaller, this presents some problems in our estimations of inbreeding depression in growth. For example, if an inbred individual is smaller than normal for its age, then the predicted age will be less than its actual age, leading to an underestimation in inbreeding depression. In reality this individual would have a shallower growth curve, as it is older than predicted. However, estimates of the asymptote should not be affected as age does not influence the trait values once individuals have finished growing. Overall, while we found convincing evidence of inbreeding depression in this wild population, we may still be underestimating its true extent due to our data collection methods, highlighting the value of such continued efforts and long-term studies.

We also used two complementary inbreeding coefficients for our analyses, yielding slightly different outcomes for the presence of inbreeding depression. While all show a negative effect of inbreeding on the trait values, only some were significant, dependent on the inbreeding coefficient used. These differences may arise from the specific variants driving inbreeding depression, because the different metrics weigh rare variants differently. For example, in tarsus length we show a significant association with *F*_ROH_ in the LMM, but no association with *F*_UniW_. This suggests that inbreeding depression in tarsus length could be caused by recessive mutations widespread in the population at intermediate or rare frequencies, which have recently come together in ROHs (Clark & Okada, 2019; Alemu et al., 2021). In contrast, for bill length *F*_ROH_ only shows an association in the LMM, whereas *F*_UniW_ shows a significant association in both the LMM and the NLMM. In this case inbreeding depression could be attributed to alleles limited to a few families within the study population. Since *F*_UniW_ gives more weight to homozygosity in rare alleles, a lower average co-ancestry results in higher scaled individual inbreeding coefficients (Zhang et al., 2022). Overall, there is no overwhelming evidence for either inbreeding coefficient being superior and is not the focus of this study. Instead, a complementary approach using multiple inbreeding coefficients may be the most enlightening, and further research is needed to fully understand how specific deleterious variants contributing to inbreeding depression can affect the coefficients and in turn the association level.

Our final aim was to determine whether particular regions of the genome contribute more (or less) to the observed inbreeding depression in our study population. In recent years, a few notable studies have had the same goal also utilising fitness data from their respective populations - Soay sheep (Stoffel et al., 2021), Hihi (Duntsch et al., 2023), and red deer (Hewett et al., 2024). Other studies have addressed the same underlying question through a fundamentally different approach, using WGS to infer loss-of-function genotypes, such as Hasselgren et al. (2024) and Lavanchy, Cumer, et al. (2024). Here, across our three traits we identified nine *sim*100kb-windows that showed a significant negative association with trait values and three with a positive association. In the cases where increased local inbreeding was positively associated with the trait, these may be regions where being homozygous for an allele is beneficial (Stoffel et al., 2021). The windows with a negative association may contain loci harbouring deleterious (recessive) alleles. Specifically, the only protein coding gene within the most highly associated window in tarsus length on chromosome 7 is COL3A1 involved in collagen synthesis, although this is partly anecdotal and non-causal. Such deleterious alleles would be expected to be kept at a lower frequency as a result of purging. In Stoffel et al. (2021), the existence of strongly deleterious genomic regions were attributed to long-term small population size and genetic drift allowing these alleles to persist. In our case - where the effective population size of the barn owl is large (Lavanchy, Cumer, et al., 2024) and extreme inbreeding is uncommon (Fig.1A) - the modest effect sizes we see of deleterious windows (Fig.3) may have prevented them from being fully purged. In addition, our equivalent analysis with local *F*_UniW_ contains no overlapping significant windows, suggesting different causal variants are being captured between metrics, or the deleterious impact is small. These analyses imply that inbreeding depression in this population likely arose from a combined effect of numerous mildly deleterious loci genome-wide, alongside a smaller number of loci with moderate or severe effects.

In conclusion, we now have a more complete picture of how inbreeding depression affects individuals throughout their early life in the wild. We find compelling evidence for inbreeding depression in three morphological traits using multiple complementing methods and show that the severity of inbreeding depression - or our ability to detect it - changes over the course of an individuals growth period. Most notably we reveal that inbreeding depression can affect weight gain at an early age. We also show that environmental impacts on traits change over the same time period which may play a role in determining the selection pressure. Finally, we show that the genetic architecture of inbreeding depression likely consists of a number of mildly deleterious (recessive) alleles genome-wide, alongside a handful of loci we identify with a more severe impact.

## 6 Data Availability

Data will be archived on acceptance. All code used is currently available at: https://github.com/annamayh/Owls/tree/418fe0c27a0114795f8e6d175d917a617581a663/Inbreedingdepression

## 7 Author Contributions

J.G., E.L. and A.M.H. conceived the study. E.L. wrote the first draft, conducted the initial analysis and calculated inbreeding coefficients. A.M.H. refined models, conducted the analysis of inbreeding effects on growth rates and genetic architecture and developed subsequent drafts. A.T., T.C. and E.L. conducted SNP-calling, phasing, mapping and quality control of the genotyped samples. A.L.R and C.S. extracted DNA and prepared samples for sequencing.

A.R. and B.A. coordinated sampling and field work. All authors contributed to revisions.

## 9 Acknowledgements

We thank the many field assistants at UNIL and Volgelwarte over the years who have collected samples and taken phenotype measurements, helping to maintain the long-term project. In particular Clara Deillon and Alexandre Roland, who we also thank for their advice. We further thank Paolo Becciu for his insights. We are grateful to all members of the Goudet group for their comments and suggestions.

