## Supplementary Fig for "Growth effects and the underlying genetic architecture of inbreeding depression in a wild raptor"

### Supplementary Figures

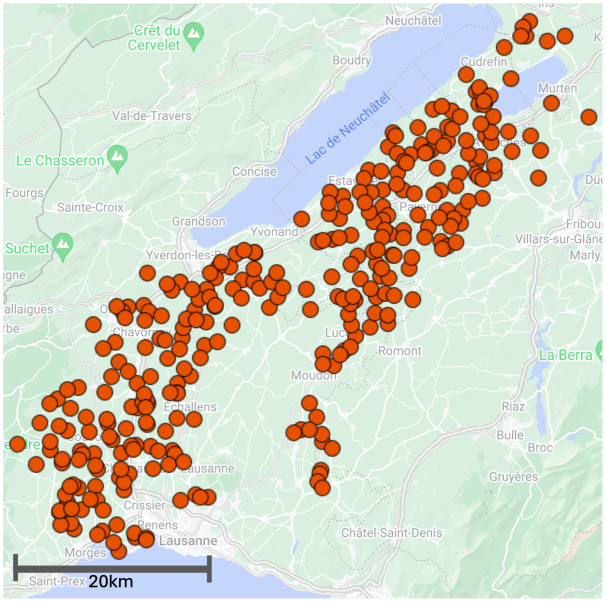


Figure 1: Nestbox locations in the study area as of 2025 field season in south west Switzerland.

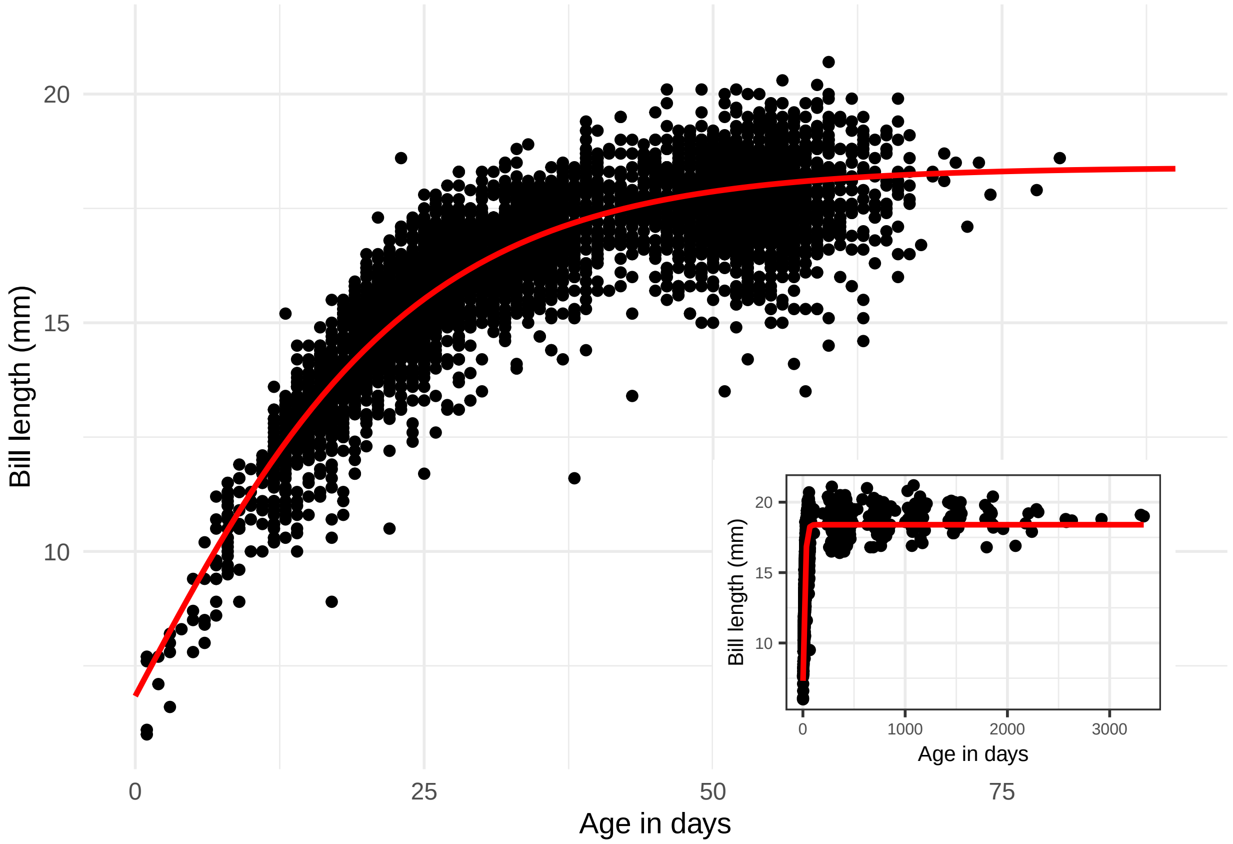


Figure 2: Gompertz growth curve used to account for age of an individual in linear mixed model, defined as: y=18.4 x exp(-0.99 x 0.932ˆx), where $x% is the age of an individual in days. Red line shows the function curve, dots are the raw data points used in the analysis. Inset plot shows the full data.

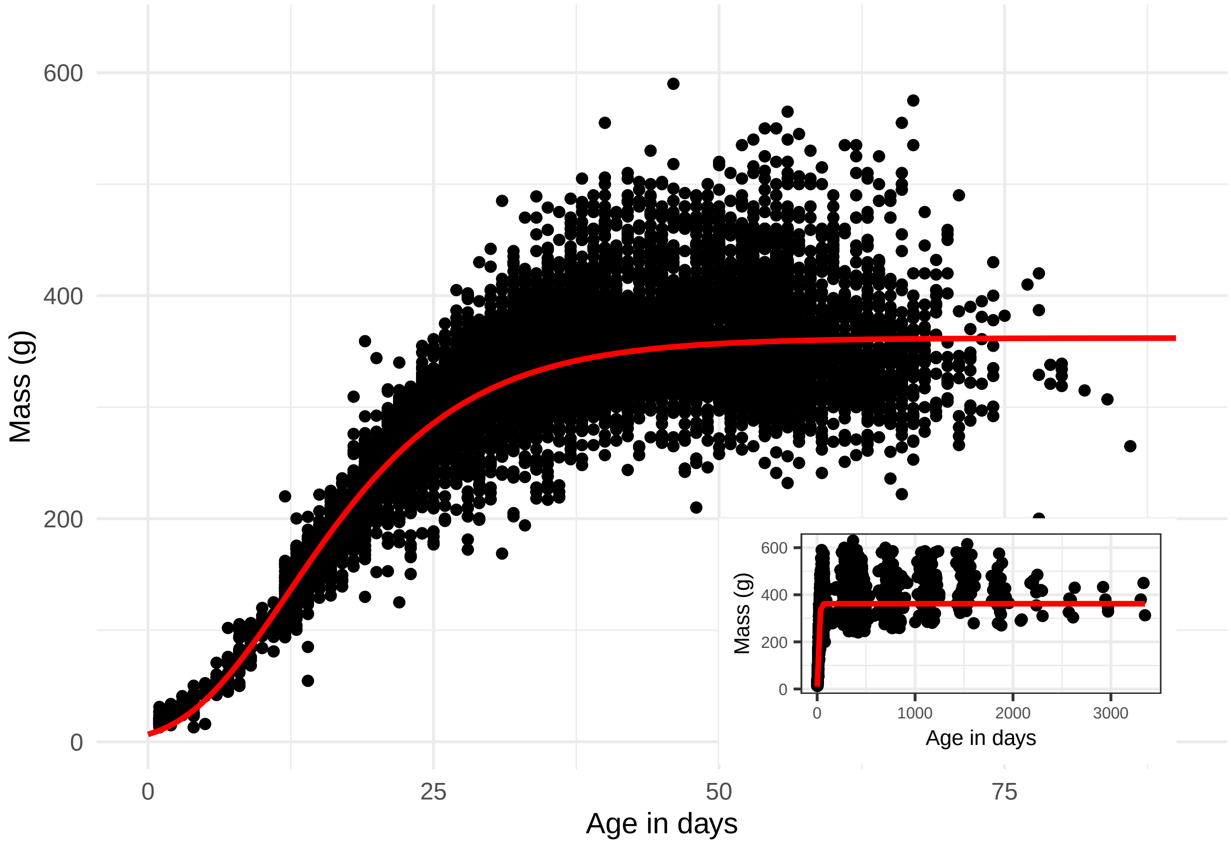


Figure 3: Gompertz growth curve used to account for age of an individual in linear mixed model, defined as: $y = 362 \cdot exp(-4.01 \cdot 0.893^x)$, where $x% is the age of an individual in days. Red line shows the function curve, dots are the raw data points used in the analysis. Inset plot shows the full data.

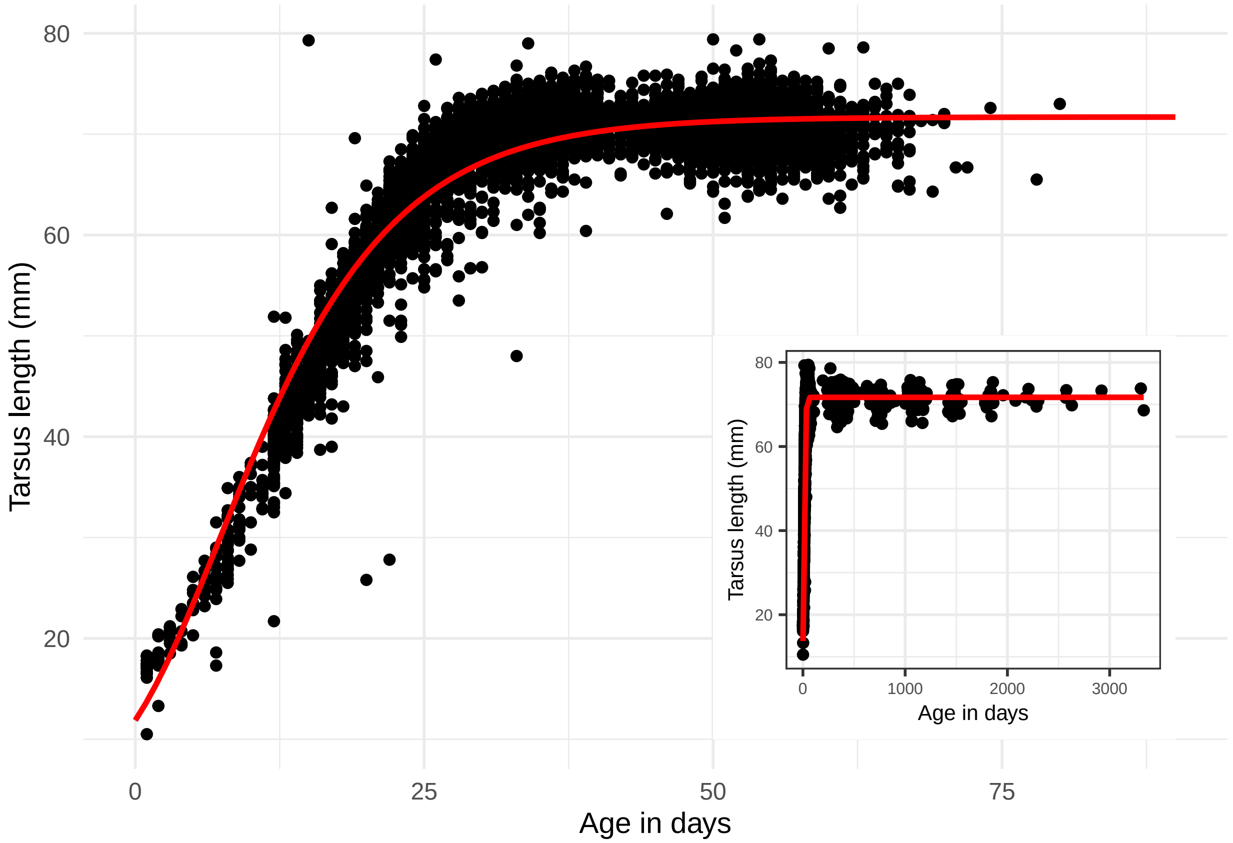


Figure 4: Gompertz growth curve used to account for age of an individual in linear mixed model, defined as: $y = 5.5 + (66.2 \cdot exp(-2.341 \cdot 0.89^{x}))$, where $x% is the age of an individual in days. Red line shows the function curve, dots are the raw data points used in the analysis. Inset plot shows the full data used.

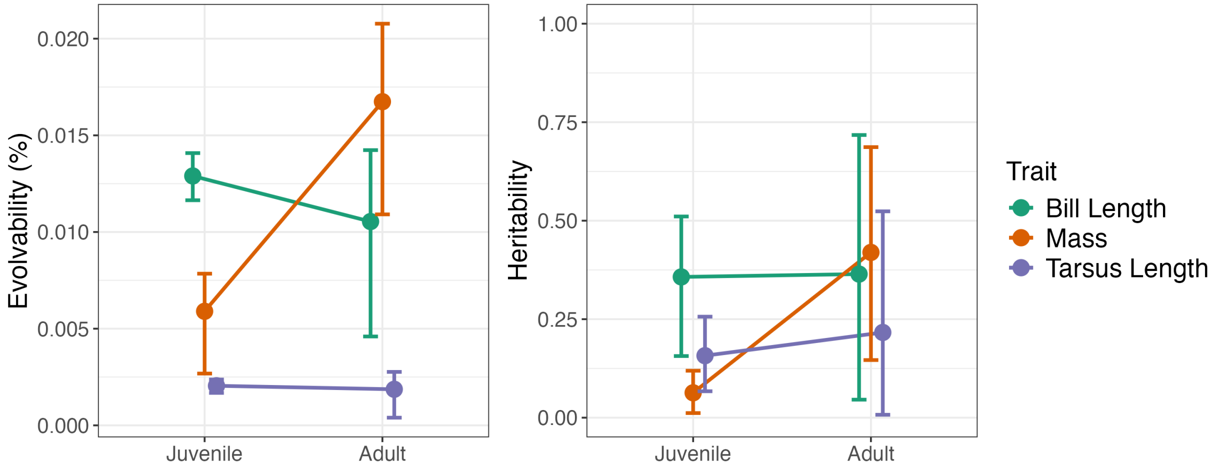


Figure 5: Estimated evolvability (additive genetic variance divided by trait mean squared, left) and narrow sense heritability (additive genetc variance divided by the total phenotypic variance, right) change between juveniles (<180 days) and adults (>180 days) for three traits with credible intervals. Calculated from the model described in section 3.2

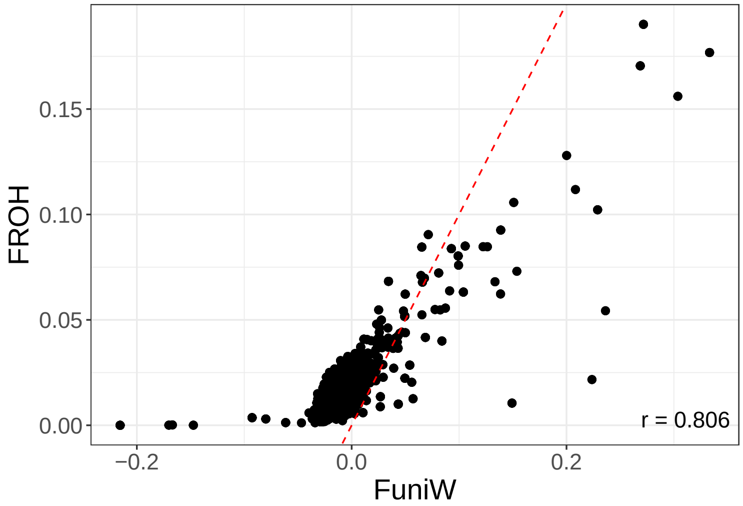


Figure 6: Correlation between two inbreeding coefficients (FROH and FuniW) within individuals. Red dashed line indicates the 1-1 line. Note that FuniW can range from -1 to 1 and FROH from 0 to 1.

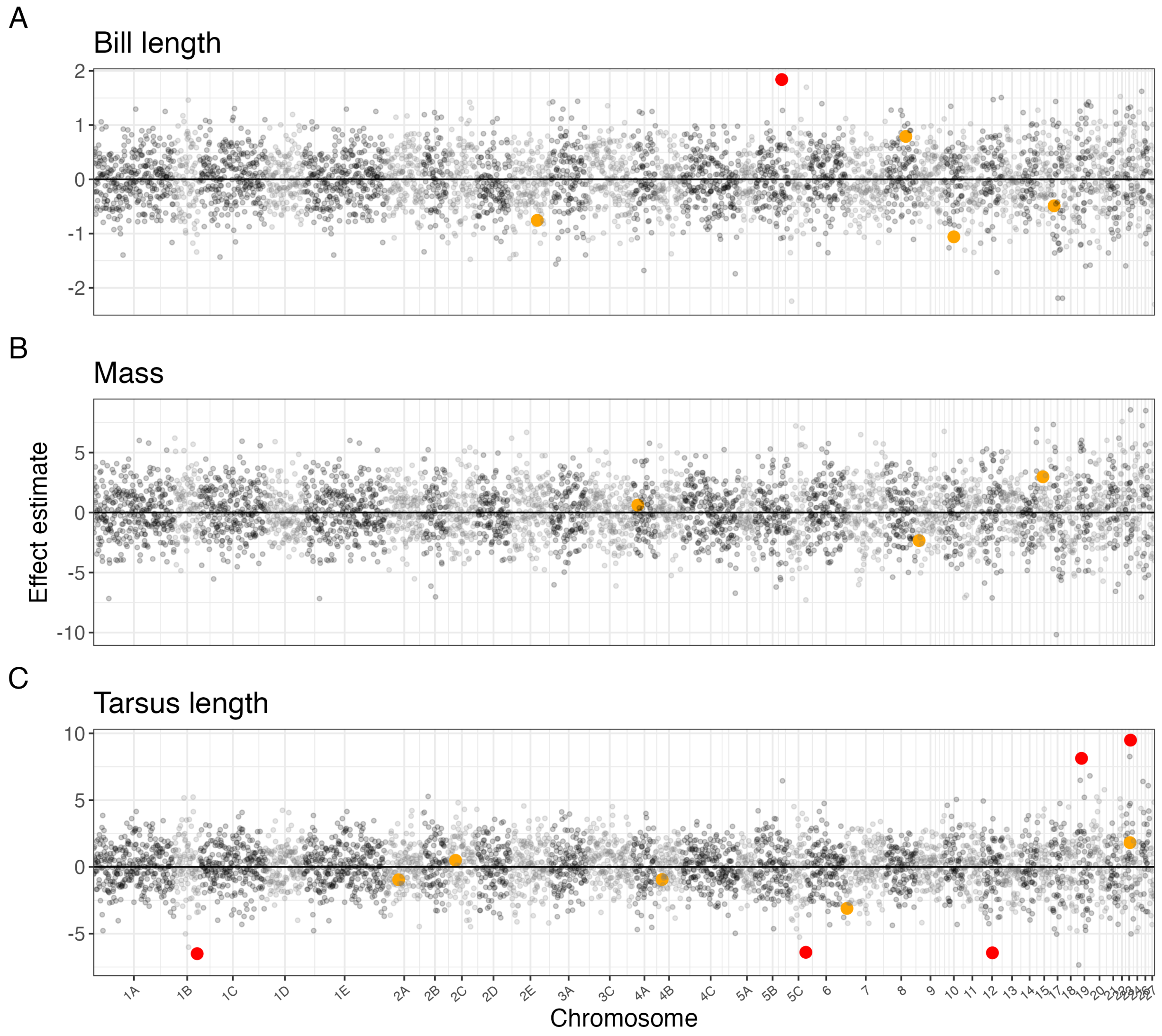


Figure 7: Estimated effect sizes of local inbreeding using FuniW in 2500 SNP windows when local FuniW and genome-wide Funi are fitted together in the same model. Red coloured dots show significant windows crossing the Bonferroni corrected significance threshold where the p-value is calculated as the number of post-warm up iterations with an opposite values to the mean estimate. Orange dots show windows that were significant using local FROH (Figure 3), no windows were significant in both approaches. Chromosome numbers are corresponding to the chicken genome based on the linkage groups identified in Topaloudis et al. (2025).

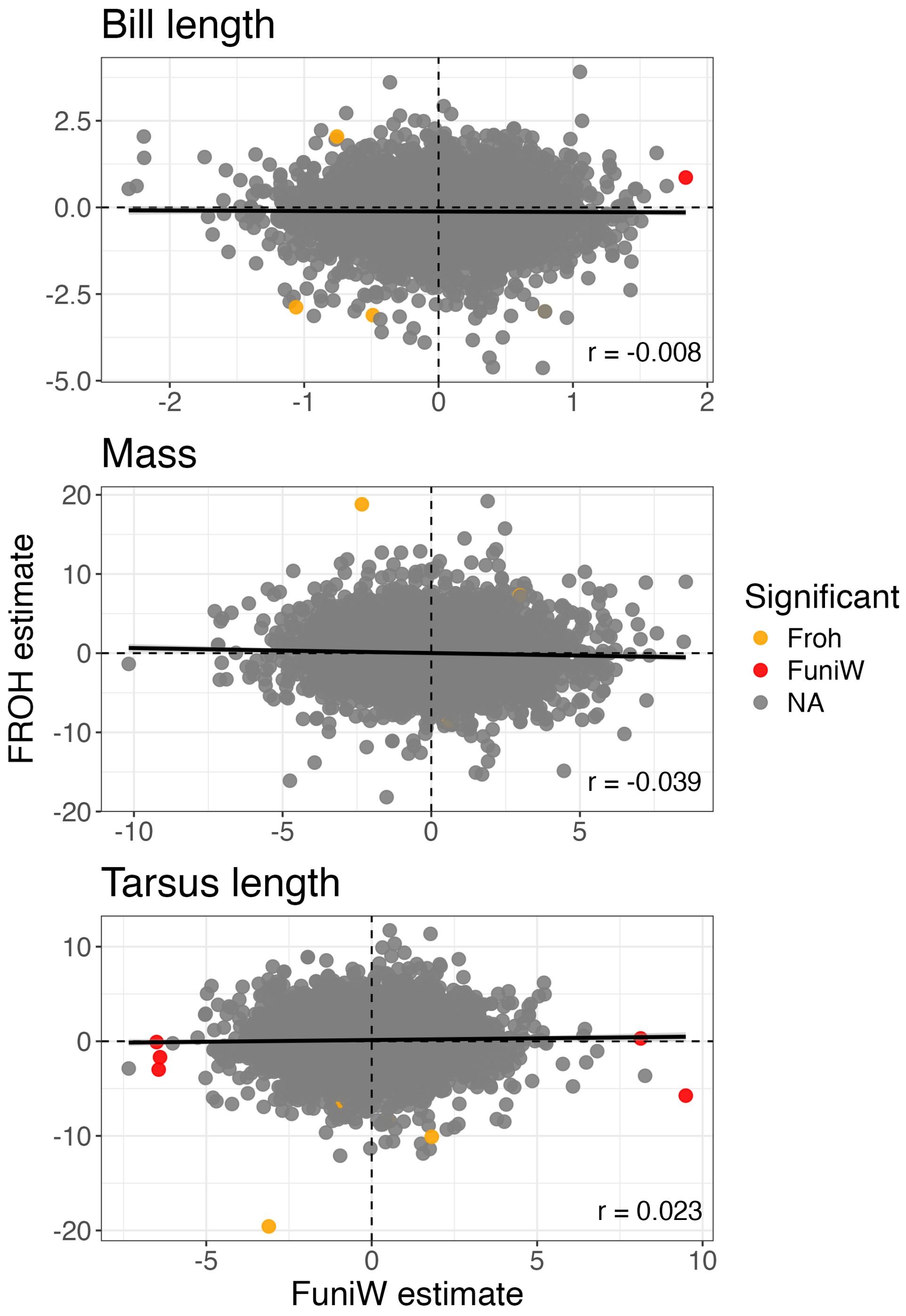


Figure 8: Correlation of estimated effect sizes within one 2500 SNP window between the two local inbreeding coefficients.
