## Supplementary Table for "Growth effects and the underlying genetic architecture of inbreeding depression in a wild raptor"

### Supplementary Methods: Model specifications and input data

Table 1: Number of unique individuals with genetic and phenotype data, with the number of records for each trait as many individuals have repeated measurements.

| Trait | Number of unique individuals | Record number |
|---|---|---|
| Bill Length | 1996 | 5596 |
| Mass | 1996 | 9001 |
| Tarsus Length | 1995 | 5731 |

#### Inbreeding depression in traits

Table 2: Model specifications for inbreeding depression in traits using a linear mixed model (LMM). All models were run with 4 MCMC chains.

| Model number | Trait | Inbreeding Coefficient | Number of iterations | Number of warm-up iterations | Thinning interval |
|---|---|---|---|---|---|
| 1.1a | BillLength | FuniW | 25000 | 7500 | 5 |
| 1.1b | BillLength | FROH | 25000 | 7500 | 5 |
| 1.2a | Mass | FuniW | 50000 | 20000 | 5 |
| 1.2b | Mass | FROH | 50000 | 20000 | 5 |
| 1.3a | LeftTarsus | FuniW | 50000 | 20000 | 5 |
| 1.3b | LeftTarsus | FROH | 50000 | 20000 | 5 |

#### Inbreeding depression in growth rates

Table 3: Model specifications for inbreeding depression in growth rates using a non-linear mixed model (NLMM). All models were run with 4 MCMC chains.

| Model number | Trait | Inbreeding Coefficient | Number of iterations | Number of warm-up iterations | Thinning interval |
|---|---|---|---|---|---|
| 2.1a | BillLength | FuniW | 35000 | 15000 | 5 |
| 2.1b | BillLength | FROH | 35000 | 15000 | 5 |
| 2.2a | Mass | FuniW | 70000 | 30000 | 5 |
| 2.2b | Mass | FROH | 70000 | 30000 | 5 |
| 2.3a | LeftTarsus | FuniW | 55000 | 15000 | 10 |
| 2.3b | LeftTarsus | FROH | 55000 | 15000 | 10 |

#### Heritability in stages

Table 4: Model specifications for heritability in stages. All models were run with 4 MCMC chains.

| Model number | Trait | Number of iterations | Number of warm-up iterations | Thinning interval |
|---|---|---|---|---|
| 3.1 | BillLength | 35000 | 5000 | 5 |
| 3.2 | Mass | 40000 | 10000 | 5 |
| 3.3 | LeftTarsus | 25000 | 8000 | 5 |

**Genetic architecture of inbreeding depression** To model the genetic architecture of inbreeding depression we used 3 chains and the default number of iterations (2000) and warm-up iterations (1000) for each window.

### Supplementary results: Inbreeding depression in traits

#### Population-level ('fixed') effects

#### Bill length

Table 5: Population-level ('fixed') effects for model 1.1.a

| | Estimate | Est.Error | Q2.5 | Q97.5 |
|---|---|---|---|---|
| Intercept | 165.94 | 1.29 | 163.51 | 168.57 |
| FuniW | -20.90 | 6.14 | -32.96 | -8.85 |
| sex2 | 3.49 | 0.35 | 2.80 | 4.19 |
| mc_age_acc | 0.97 | 0.01 | 0.96 | 0.98 |
| rank | -0.25 | 0.09 | -0.44 | -0.07 |

**Note:** [a]Est.error, Q2.5 and Q97.5 show the error, lower and upper confidence intervals around the estimate all on the trait scale: mm/0.1. FuniW indicates the estimated effect of the inbreeding coefficient. Sex2 shows the effect of being female compared to male (reference), mc_age_acc indicates the mean centred variable to account for the age of an individual, so all estimates are for an individual of an average age. See methods for more details on accounting for age. Rank is the ordered number that an individual was born in the nest i.e. individual born first is given a number 1 and so on.

Table 6: Population-level ('fixed') effects for model 1.1.b

| | Estimate | Est.Error | Q2.5 | Q97.5 |
|---|---|---|---|---|
| Intercept | 166.71 | 1.31 | 164.22 | 169.38 |
| FROH | -35.13 | 12.15 | -59.17 | -11.43 |
| sex2 | 3.43 | 0.36 | 2.73 | 4.13 |
| mc_age_acc | 0.97 | 0.01 | 0.96 | 0.98 |
| rank | -0.25 | 0.09 | -0.44 | -0.07 |

**Note:** [a]Est.error, Q2.5 and Q97.5 show the error, lower and upper confidence intervals around the estimate all on the trait scale: mm/0.1. FROH indicates the estimated effect of the inbreeding coefficient. Sex2 shows the effect of being female compared to male (reference), mc_age_acc indicates the mean centred variable to account for the age of an individual, so all estimates are for an individual of an average age. See methods for more details on accounting for age. Rank is the ordered number that an individual was born in the nest i.e. individual born first is given a number 1 and so on.

**Mass**

Table 7: Population-level ('fixed') for model 1.2.a

| | Estimate | Est.Error | Q2.5 | Q97.5 |
|---|---|---|---|---|
| Intercept | 337.58 | 7.25 | 323.11 | 351.58 |
| FuniW | -27.74 | 29.05 | -85.09 | 29.05 |
| sex2 | 16.05 | 1.30 | 13.51 | 18.61 |
| rank | -0.95 | 0.40 | -1.75 | -0.16 |
| mc_age_acc | 1.06 | 0.01 | 1.05 | 1.07 |

**Note:** [a]Est.error, Q2.5 and Q97.5 show the error, lower and upper confidence intervals around the estimate all on the trait scale: grams. FuniW indicates the estimated effect of the inbreeding coefficient. Sex2 shows the effect of being female compared to male (reference), mc_age_acc indicates the mean centred variable to account for the age of an individual, so all estimates are for an individual of an average age. See methods for more details on accounting for age. Rank is the ordered number that an individual was born in the nest i.e. individual born first is given a number 1 and so on.

Table 8: Population-level ('fixed') for model 1.2.b

| | Estimate | Est.Error | Q2.5 | Q97.5 |
|---|---|---|---|---|
| Intercept | 338.94 | 7.29 | 324.49 | 353.41 |
| FROH | -61.16 | 56.74 | -171.25 | 50.51 |
| sex2 | 15.94 | 1.29 | 13.40 | 18.45 |
| rank | -0.95 | 0.40 | -1.74 | -0.15 |
| mc_age_acc | 1.06 | 0.01 | 1.04 | 1.07 |

**Note:** [a]Est.error, Q2.5 and Q97.5 show the error, lower and upper confidence intervals around the estimate all on the trait scale grams. FROH indicates the estimated effect of the inbreeding coefficient. Sex2 shows the effect of being female compared to male (reference), mc_age_acc indicates the mean centred variable to account for the age of an individual, so all estimates are for an individual of an average age. See methods for more details on accounting for age. Rank is the ordered number that an individual was born in the nest i.e. individual born first is given a number 1 and so on.

**Tarsus length**

Table 9: Population-level ('fixed') for model 1.3.a

| | Estimate | Est.Error | Q2.5 | Q97.5 |
|---|---|---|---|---|
| Intercept | 670.11 | 3.08 | 663.65 | 675.87 |
| FuniW | -34.80 | 21.61 | -77.40 | 7.44 |
| sex2 | -4.75 | 1.11 | -6.94 | -2.58 |
| rank | -6.19 | 0.32 | -6.81 | -5.54 |
| mc_age_acc | 1.03 | 0.00 | 1.03 | 1.04 |

**Note:** [a]Est.error, Q2.5 and Q97.5 show the error, lower and upper confidence intervals around the estimate all on the trait scale mm/0.1. FuniW indicates the estimated effect of the inbreeding coefficient. Sex2 shows the effect of being female compared to male (reference), mc_age_acc indicates the mean centred variable to account for the age of an individual, so all estimates are for an individual of an average age. See methods for more details accounting for age. Rank is the ordered number that an individual was born in the nest i.e. individual born first is given a number 1 and so on.

Table 10: Population-level ('fixed') for model 1.3.b

| | Estimate | Est.Error | Q2.5 | Q97.5 |
|---|---|---|---|---|
| Intercept | 672.28 | 3.21 | 665.65 | 678.22 |
| FROH | -108.31 | 43.40 | -193.67 | -22.58 |
| sex2 | -4.96 | 1.10 | -7.12 | -2.82 |
| rank | -6.20 | 0.33 | -6.84 | -5.56 |
| mc_age_acc | 1.03 | 0.00 | 1.03 | 1.04 |

**Note:** [a]Est.error, Q2.5 and Q97.5 show the error, lower and upper confidence intervals around the estimate all on the trait scale mm/0.1. FROH indicates the estimated effect of the inbreeding coefficient. Sex2 shows the effect of being female compared to male (reference), mc_age_acc indicates the mean centred variable to account for the age of an individual, so all estimates are for an individual of an average age. See methods for more details accounting for age. Rank is the ordered number that an individual was born in the nest i.e. individual born first is given a number 1 and so on.

#### Group level ('Random') effects

#### Bill length

Table 11: Group level ('random') effects for model 1.1.a

| group | estimate | std.error | conf.low | conf.high | N levels |
|---|---|---|---|---|---|
| clutch_merge | 2.17 | 0.32 | 1.50 | 2.74 | 699 |
| month | 1.84 | 0.99 | 0.69 | 4.46 | 7 |
| nestboxID | 0.93 | 0.45 | 0.07 | 1.76 | 303 |
| Observer | 2.48 | 0.42 | 1.76 | 3.40 | 46 |
| RingId | 3.36 | 0.16 | 3.03 | 3.67 | 1996 |
| RingId_pe | 0.53 | 0.38 | 0.02 | 1.42 | 1996 |
| year | 2.19 | 0.69 | 1.11 | 3.78 | 24 |
| Residual | 6.29 | 0.07 | 6.15 | 6.43 | NA |

**Note:** [a]Each group shows a random effect fitted in the model with estimates, error and confidence intervals for all groups in the model. N levels indicates the number of different levels for each group. Clutch_merge is the clutch identity that and individual was born and raised (see methods for explanation). Month and year are those in which an individual is born. NestbooxID is the individual nestbox location identity. RingId is the additive genetic variance (Va) and RingId_pe is the permanent environment of the individual to account for repeated measurements. Observer is the individual taking measurements.

Table 12: Group level ('random') effects for model 1.1.b

| group | estimate | std.error | conf.low | conf.high | N levels |
|---|---|---|---|---|---|
| clutch_merge | 2.17 | 0.33 | 1.49 | 2.77 | 699 |
| month | 1.85 | 1.03 | 0.69 | 4.46 | 7 |
| nestboxID | 0.98 | 0.45 | 0.08 | 1.80 | 303 |
| Observer | 2.48 | 0.42 | 1.76 | 3.43 | 46 |
| RingId | 3.36 | 0.17 | 3.02 | 3.68 | 1996 |
| RingId_pe | 0.54 | 0.39 | 0.02 | 1.41 | 1996 |
| year | 2.20 | 0.70 | 1.11 | 3.85 | 24 |
| Residual | 6.29 | 0.07 | 6.15 | 6.44 | NA |

**Note:** [a]Each group shows a random effect fitted in the model with estimates, error and confidence intervals for all groups in the model. N levels indicates the number of different levels for each group. Clutch_merge is the clutch identity that and individual was born and raised (see methods for explanation). Month and year are those in which an individual is born. NestbooxID is the individual nestbox location identity. RingId is the additive genetic variance (Va) and RingId_pe is the permanent environment of the individual to account for repeated measurements. Observer is the individual taking measurements.

**Mass**

Table 13: Group level ('random') effects for model 1.2.a

| group | estimate | std.error | conf.low | conf.high | N levels |
|---|---|---|---|---|---|
| clutch_merge | 15.46 | 0.96 | 13.55 | 17.33 | 699 |
| month | 7.84 | 4.11 | 2.97 | 18.61 | 7 |
| nestboxID | 2.70 | 1.73 | 0.13 | 6.34 | 303 |
| Observer | 39.13 | 4.46 | 31.31 | 48.83 | 62 |
| RingId | 8.67 | 2.07 | 3.23 | 11.44 | 1996 |
| RingId_pe | 5.43 | 2.97 | 0.30 | 10.71 | 1996 |
| year | 9.94 | 2.49 | 6.05 | 15.72 | 24 |
| Residual | 37.96 | 0.31 | 37.35 | 38.58 | NA |

**Note:** [a]Each group shows a random effect fitted in the model with estimates, error and confidence intervals for all groups in the model. N levels indicates the number of different levels for each group. Clutch_merge is the clutch identity that and individual was born and raised (see methods for explanation). Month and year are those in which an individual is born. NestbooxID is the individual nestbox location identity. RingId is the additive genetic variance (Va) and RingId_pe is the permanent environment of the individual to account for repeated measurements. Observer is the individual taking measurements.

Table 14: Group level ('random') effects for model 1.2.b

| group | estimate | std.error | conf.low | conf.high | N levels |
|---|---|---|---|---|---|
| clutch_merge | 15.46 | 0.96 | 13.58 | 17.33 | 699 |
| month | 7.93 | 4.21 | 3.00 | 18.81 | 7 |
| nestboxID | 2.70 | 1.72 | 0.13 | 6.33 | 303 |
| Observer | 39.22 | 4.46 | 31.36 | 48.78 | 62 |
| RingId | 8.79 | 1.99 | 3.72 | 11.43 | 1996 |
| RingId_pe | 5.26 | 2.95 | 0.28 | 10.56 | 1996 |
| year | 9.96 | 2.49 | 6.07 | 15.66 | 24 |

| group | estimate | std.error | conf.low | conf.high | N levels |
|---|---|---|---|---|---|
| Residual | 37.96 | 0.31 | 37.35 | 38.57 | NA |

**Note:** [a]Each group shows a random effect fitted in the model with estimates, error and confidence intervals for all groups in the model. N levels indicates the number of different levels for each group. Clutch_merge is the clutch identity that and individual was born and raised (see methods for explanation). Month and year are those in which an individual is born. NestbooxID is the individual nestbox location identity. RingId is the additive genetic variance (Va) and RingId_pe is the permanent environment of the individual to account for repeated measurements. Observer is the individual taking measurements.

#### Tarsus length

Table 15: Group level ('random') effects for model 1.3.a

| group | estimate | std.error | conf.low | conf.high | N levels |
|---|---|---|---|---|---|
| clutch_merge | 10.10 | 0.88 | 8.35 | 11.78 | 699 |
| month | 3.41 | 3.01 | 0.22 | 11.34 | 7 |
| nestboxID | 3.48 | 1.46 | 0.37 | 6.04 | 303 |
| Observer | 5.13 | 1.07 | 3.32 | 7.48 | 48 |
| RingId | 9.51 | 0.64 | 8.18 | 10.71 | 1995 |
| RingId_pe | 1.68 | 1.25 | 0.06 | 4.61 | 1995 |
| year | 6.16 | 1.92 | 3.21 | 10.67 | 24 |
| Residual | 23.39 | 0.26 | 22.88 | 23.93 | NA |

**Note:** [a]Each group shows a random effect fitted in the model with estimates, error and confidence intervals for all groups in the model. N levels indicates the number of different levels for each group. Clutch_merge is the clutch identity that and individual was born and raised (see methods for explanation). Month and year are those in which an individual is born. NestbooxID is the individual nestbox location identity. RingId is the additive genetic variance (Va) and RingId_pe is the permanent environment of the individual to account for repeated measurements. Observer is the individual taking measurements.

Table 16: Group level ('random') effects for model 1.3.b

| group | estimate | std.error | conf.low | conf.high | N levels |
|---|---|---|---|---|---|
| clutch_merge | 10.18 | 0.87 | 8.44 | 11.85 | 699 |
| month | 3.41 | 2.97 | 0.20 | 11.11 | 7 |
| nestboxID | 3.43 | 1.48 | 0.36 | 6.03 | 303 |
| Observer | 5.11 | 1.06 | 3.32 | 7.45 | 48 |
| RingId | 9.44 | 0.65 | 8.10 | 10.64 | 1995 |
| RingId_pe | 1.74 | 1.28 | 0.07 | 4.74 | 1995 |
| year | 6.25 | 1.96 | 3.24 | 10.80 | 24 |
| Residual | 23.39 | 0.26 | 22.88 | 23.91 | NA |

**Note:** [a]Each group shows a random effect fitted in the model with estimates, error and confidence intervals for all groups in the model. N levels indicates the number of different levels for each group. Clutch_merge is the clutch identity that and individual was born and raised (see methods for explanation). Month and year are those in which an individual is born. NestbooxID is the individual nestbox location identity. RingId is the additive genetic variance (Va) and RingId_pe is the permanent environment of the individual to account for repeated measurements. Observer is the individual taking measurements.

### Supplementary Results: Inbreeding depression in growth rates

#### Bill length

Table 17: Population-level ('fixed') effects for model 2.1a, with bill length as the response variable. All estimates are on the trait scale: mm*0.1. Estimates are for each parameter, where asym is the asymptote, b is the initiation of the growth curve and c is the growth slope.

| | Estimate | Est.Error | Q2.5 | Q97.5 |
|---|---|---|---|---|
| asym__Intercept | 182.50 | 1.47 | 179.62 | 185.42 |
| asym__FuniW | -17.70 | 6.61 | -30.63 | -4.68 |
| asym__rank | -0.39 | 0.14 | -0.67 | -0.12 |
| asym__sex2 | 4.14 | 0.38 | 3.39 | 4.88 |
| b__Intercept | 1.05 | 0.02 | 1.02 | 1.08 |
| c__Intercept | -0.08 | 0.00 | -0.08 | -0.07 |
| c__FuniW | -0.01 | 0.01 | -0.04 | 0.02 |
| c__rank | 0.00 | 0.00 | 0.00 | 0.00 |
| c__sex2 | 0.00 | 0.00 | 0.00 | 0.00 |

Table 18: Population-level ('fixed') effects for model 2.1a, with bill length as the response variable. All estimates are on the trait scale: mm*0.1. Estimates are for each parameter, where asym is the asymptote, b is the initiation of the growth curve and c is the growth slope.

| | Estimate | Est.Error | Q2.5 | Q97.5 |
|---|---|---|---|---|
| asym__Intercept | 182.91 | 1.47 | 180.02 | 185.88 |
| asym__FROH | -14.44 | 8.77 | -31.52 | 2.55 |
| asym__rank | -0.39 | 0.14 | -0.67 | -0.12 |
| asym__sex2 | 4.15 | 0.38 | 3.40 | 4.90 |
| b__Intercept | 1.05 | 0.02 | 1.02 | 1.08 |
| c__Intercept | -0.08 | 0.00 | -0.08 | -0.07 |
| c__FROH | 0.00 | 0.03 | -0.05 | 0.05 |
| c__rank | 0.00 | 0.00 | 0.00 | 0.00 |
| c__sex2 | 0.00 | 0.00 | 0.00 | 0.00 |

#### Mass

Table 19: Population-level ('fixed') effects for model 2.2a, with mass as the response variable. All estimates are on the trait scale: grams. Estimates are for each parameter, where asym is the asymptote, b is the initiation of the growth curve and c is the growth slope.

| | Estimate | Est.Error | Q2.5 | Q97.5 |
|---|---|---|---|---|
| asym__Intercept | 373.90 | 7.06 | 359.97 | 387.55 |
| asym__FuniW | 0.64 | 17.71 | -34.40 | 35.19 |
| asym__rank | 2.57 | 0.57 | 1.46 | 3.67 |

| | Estimate | Est.Error | Q2.5 | Q97.5 |
|---|---|---|---|---|
| asym_sex2 | 23.87 | 1.51 | 20.92 | 26.82 |
| b_Intercept | 3.72 | 0.09 | 3.55 | 3.90 |
| c_Intercept | 0.89 | 0.00 | 0.89 | 0.90 |
| c_FuniW | 0.03 | 0.01 | 0.01 | 0.06 |
| c_rank | 0.00 | 0.00 | 0.00 | 0.00 |
| c_sex2 | 0.00 | 0.00 | 0.00 | 0.01 |

Table 20: Population-level ('fixed') effects for model 2.2a, with mass as the response variable. All estimates are on the trait scale: grams. Estimates are for each parameter, where asym is the asymptote, b is the initiation of the growth curve and c is the growth slope.

| | Estimate | Est.Error | Q2.5 | Q97.5 |
|---|---|---|---|---|
| asym_Intercept | 374.07 | 7.04 | 360.03 | 387.88 |
| asym_FROH | 1.79 | 19.33 | -36.04 | 40.02 |
| asym_rank | 2.56 | 0.57 | 1.44 | 3.69 |
| asym_sex2 | 23.84 | 1.49 | 20.91 | 26.74 |
| b_Intercept | 3.72 | 0.09 | 3.55 | 3.90 |
| c_Intercept | 0.89 | 0.00 | 0.89 | 0.89 |
| c_FROH | 0.08 | 0.03 | 0.02 | 0.13 |
| c_rank | 0.00 | 0.00 | 0.00 | 0.00 |
| c_sex2 | 0.00 | 0.00 | 0.00 | 0.01 |

**Tarsus length**

Table 21: Population-level ('fixed') effects for model 2.3a, with tarsus length as the response variable. All estimates are on the trait scale: mm*0.1. Estimates are for each parameter, where asym is the asymptote, b is the initiation of the growth curve and c is the growth slope.

| | Estimate | Est.Error | Q2.5 | Q97.5 |
|---|---|---|---|---|
| asym1_Intercept | 142.83 | 3.76 | 135.32 | 150.14 |
| asym_Intercept | 580.75 | 5.41 | 571.16 | 591.54 |
| asym_FuniW | -26.94 | 14.73 | -55.87 | 2.23 |
| asym_rank | -4.28 | 0.33 | -4.93 | -3.62 |
| asym_sex2 | -3.57 | 0.89 | -5.33 | -1.86 |
| b_Intercept | 4.80 | 0.14 | 4.53 | 5.07 |
| c_Intercept | 0.86 | 0.00 | 0.86 | 0.86 |
| c_FuniW | -0.02 | 0.01 | -0.04 | 0.01 |
| c_rank | 0.00 | 0.00 | 0.00 | 0.00 |
| c_sex2 | 0.00 | 0.00 | 0.00 | 0.00 |

Table 22: Population-level ('fixed') effects for model 2.3a, with tarsus length as the response variable. All estimates are on the trait scale: mm*0.1. Estimates are for each parameter, where asym is the asymptote, b is the initiation of the growth curve and c is the growth slope.

| | Estimate | Est.Error | Q2.5 | Q97.5 |
|---|---|---|---|---|
| asym1_Intercept | 142.84 | 3.75 | 135.26 | 150.16 |
| asym_Intercept | 581.49 | 5.33 | 571.87 | 592.28 |
| asym_FROH | -32.09 | 18.41 | -68.37 | 4.11 |
| asym_rank | -4.28 | 0.33 | -4.93 | -3.64 |
| asym_sex2 | -3.61 | 0.88 | -5.34 | -1.90 |
| b_Intercept | 4.80 | 0.14 | 4.53 | 5.07 |
| c_Intercept | 0.86 | 0.00 | 0.86 | 0.86 |
| c_FROH | -0.02 | 0.02 | -0.06 | 0.03 |
| c_rank | 0.00 | 0.00 | 0.00 | 0.00 |
| c_sex2 | 0.00 | 0.00 | 0.00 | 0.00 |

### Supplementary Results: Heritability in stages

#### Bill length

Table 23: Proportion of trait variance explained with upper and lower 95% credible intervals across juvenile and adult stages in bill length (Model 3.1)

| Variance component | Mean | Lwr | Upr | Growth stage |
|---|---:|---:|---:|---|
| Va | 0.3572636 | 0.1564421 | 0.5106998 | Juvenile |
| pe | 0.0119251 | 0.0000123 | 0.0582888 | Juvenile |
| year | 0.1503636 | 0.0330302 | 0.3744343 | Juvenile |
| month | 0.1972449 | 0.0223294 | 0.6467282 | Juvenile |
| nestbox | 0.0484499 | 0.0009847 | 0.1229567 | Juvenile |
| clutch | 0.1186463 | 0.0294543 | 0.2235829 | Juvenile |
| rank | 0.0159562 | 0.0000110 | 0.1048970 | Juvenile |
| resid | 0.1001504 | 0.0448105 | 0.1378002 | Juvenile |
| Va | 0.3644800 | 0.0457307 | 0.7174967 | Adult |
| pe | 0.0884251 | 0.0001041 | 0.3879642 | Adult |
| year | 0.0349542 | 0.0000477 | 0.1608000 | Adult |
| month | 0.1164196 | 0.0001273 | 0.5999059 | Adult |
| nestbox | 0.0879459 | 0.0002079 | 0.3024693 | Adult |
| clutch | 0.1664548 | 0.0004991 | 0.4982527 | Adult |
| rank | 0.0787613 | 0.0000404 | 0.5106715 | Adult |
| resid | 0.0625591 | 0.0247871 | 0.0984602 | Adult |

**Note:** [a]Va = additive genetic variance, pe = permanent environment, year = birth year of individual, month = birth month of individual, nestbox = nextbox identity number, clutch = clutch identity, rank = order individual was born in the clutch, resid = residual variance

#### Mass

Table 24: Proportion of trait variance explained with upper and lower 95% credible intervals across juvenile and adult stages in mass (Model 3.2)

| Variance component | Mean | Lwr | Upr | Growth stage |
|---|---:|---:|---:|---|
| Va | 0.0631515 | 0.0117135 | 0.1191721 | Juvenile |
| pe | 0.0133380 | 0.0000153 | 0.0603795 | Juvenile |
| year | 0.4252267 | 0.2305593 | 0.6394049 | Juvenile |
| month | 0.0769418 | 0.0096666 | 0.3002773 | Juvenile |
| nestbox | 0.0351652 | 0.0003056 | 0.0962534 | Juvenile |
| clutch | 0.3515340 | 0.2035100 | 0.5045833 | Juvenile |
| rank | 0.0142539 | 0.0003170 | 0.0686614 | Juvenile |
| resid | 0.0203891 | 0.0124060 | 0.0280163 | Juvenile |
| Va | 0.4194884 | 0.1462488 | 0.6867477 | Adult |
| pe | 0.0724692 | 0.0000741 | 0.3293871 | Adult |
| year | 0.3300758 | 0.1022045 | 0.5897525 | Adult |
| month | 0.0353634 | 0.0000180 | 0.2584701 | Adult |
| nestbox | 0.0348361 | 0.0000350 | 0.1648753 | Adult |
| clutch | 0.0639680 | 0.0001021 | 0.2636568 | Adult |

| Variance component | Mean | Lwr | Upr | Growth stage |
|---|---|---|---|---|
| rank | 0.0355673 | 0.0000192 | 0.2399476 | Adult |
| resid | 0.0082317 | 0.0046815 | 0.0124416 | Adult |

**Note:** [a]Va = additive genetic variance, pe = permanent environment, year = birth year of individual, month = birth month of individual, nestbox = nextbox identity number, clutch = clutch identity, rank = order individual was born in the clutch, resid = residual variance

**Tarsus length**

Table 25: Proportion of trait variance explained with upper and lower 95% credible intervals across juvenile and adult stages in tarsus length (Model 3.3)

| Variance component | Mean | Lwr | Upr | Growth stage |
|---|---|---|---|---|
| Va | 0.1573628 | 0.0670568 | 0.2563833 | Juvenile |
| pe | 0.0111148 | 0.0000126 | 0.0541081 | Juvenile |
| year | 0.1087639 | 0.0232208 | 0.2812778 | Juvenile |
| month | 0.0481484 | 0.0001953 | 0.2571111 | Juvenile |
| nestbox | 0.0309919 | 0.0002933 | 0.0869597 | Juvenile |
| clutch | 0.2358616 | 0.1050796 | 0.3723033 | Juvenile |
| rank | 0.3865386 | 0.1622405 | 0.7052034 | Juvenile |
| resid | 0.0212180 | 0.0100817 | 0.0311292 | Juvenile |
| Va | 0.2162974 | 0.0073922 | 0.5237505 | Adult |
| pe | 0.0814205 | 0.0001039 | 0.3430716 | Adult |
| year | 0.0428162 | 0.0003077 | 0.1460080 | Adult |
| month | 0.1516185 | 0.0084719 | 0.5284043 | Adult |
| nestbox | 0.1555879 | 0.0013896 | 0.3849457 | Adult |
| clutch | 0.1618121 | 0.0007521 | 0.4472950 | Adult |
| rank | 0.1798019 | 0.0270076 | 0.5286926 | Adult |
| resid | 0.0106454 | 0.0048906 | 0.0162835 | Adult |

**Note:** [a]Va = additive genetic variance, pe = permanent environment, year = birth year of individual, month = birth month of individual, nestbox = nextbox identity number, clutch = clutch identity, rank = order individual was born in the clutch, resid = residual variance

### Supplementary Results: Genetic architecture of inbreeding depression

#### Correlation of genome-wide and window inbreeding coefficients within individuals

Table 26: Correlation Table of genome wide and window inbreeding coefficients for 2 inbreeding coefficients.

| | Genome-wide FHBD | Window FHBD | Genome-wide FuniW | Window FuniW |
|---|---|---|---|---|
| Genome-wide FHBD | 1.000 | NA | NA | NA |
| Window FHBD | 0.104 | 1.000 | NA | NA |
| Genome-wide FuniW | 0.831 | 0.108 | 1.000 | NA |
| Window FuniW | 0.001 | -0.001 | 0.001 | 1 |

Table 27: Genetic architecture of inbreeding depression in bill length: Table shows the few 2500-SNP windows that cross the bonferonni corrected signifiance threshold

| Chromosome | Window | estimate | lwr__CI | upr__CI | p__val |
|---|---|---|---|---|---|
| 2E | pr__hbd__super__scaffold__9__wind__109 | 2.04 | 1.11 | 2.97 | 1e-14 |
| 8 | pr__hbd__super__scaffold__1000006__wind__78 | -3.00 | -4.49 | -1.53 | 1e-14 |
| 10 | pr__hbd__super__scaffold__41__wind__37 | -2.88 | -4.30 | -1.51 | 1e-14 |
| 17 | pr__hbd__super__scaffold__11__wind__13 | -3.11 | -4.96 | -1.29 | 1e-14 |

[a] Chromosome number is based on the synteny with the chicken genome, window is a descriptive variable for locating the position within the chromosome, using the super scaffold number and window number given that each non-overlapping sliding window is 2500-SNPs. Estimate, lwr__CI and upr__CI is the estimated effect with lower and upper credible intervals of being inbred (F=1) at a window on bill length on the trait scale of mm/0.1. P-values were calculated as the proportion of post-warm up draws that were of the opposite value to the mean estimate of the effect of inbreeding

Table 28: Genetic architecture of inbreeding depression in mass: Table shows the few 2500-SNP windows that cross the bonferonni corrected signifiance threshold.

| Chromosome | Window | estimate | lwr__CI | upr__CI | p__val |
|---|---|---|---|---|---|
| 4A | pr__hbd__super__scaffold__5__wind__18 | -8.51 | -13.41 | -3.66 | 3.333333e-14 |
| 9 | pr__hbd__super__scaffold__10__wind__11 | 18.80 | 8.17 | 29.04 | 3.333333e-14 |
| 15 | pr__hbd__super__scaffold__44__wind__22 | 7.30 | 3.34 | 11.21 | 3.333333e-14 |

[a] Chromosome number is based on the synteny with the chicken genome, window is a descriptive variable for locating the position within the chromosome, using the super scaffold number and window number given that each non-overlapping sliding window is 2500-SNPs. Estimate, lwr__CI and upr__CI is the estimated effect with lower and upper credible intervals of being inbred (F=1) at a window on mass on the trait scale in grams. P-values were calculated as the proportion of post-warm up draws that were of the opposite value to the mean estimate of the effect of inbreeding

Table 29: Genetic architecture of inbreeding depression in tarsus length: Table shows the few 2500-SNP windows that cross the bonferonni corrected signifiance threshold.

| Chromosome | Window | estimate | lwr_CI | upr_CI | p_val |
|---|---|---|---|---|---|
| 2A | pr_hbd_super_scaffold_38_wind_54 | -7.29 | -11.74 | -2.69 | 3.333333e-14 |
| 2C | pr_hbd_super_scaffold_48_wind_36 | -8.35 | -13.40 | -3.13 | 3.333333e-14 |
| 4B | pr_hbd_super_scaffold_23_wind_27 | -6.27 | -10.21 | -2.39 | 3.333333e-14 |
| 7 | pr_hbd_super_scaffold_7_wind_7 | -19.57 | -28.90 | -9.70 | 3.333333e-14 |
| 23 | pr_hbd_super_scaffold_20_wind_20 | -10.10 | -15.66 | -4.50 | 3.333333e-14 |

[a] Chromosome number is based on the synteny with the chicken genome, window is a descriptive variable for locating the position within the chromosome, using the super scaffold number and window number given that each non-overlapping sliding window is 2500-SNPs. Estimate, lwr_CI and upr_CI is the estimated effect with lower and upper credible intervals of being inbred (F=1) at a window on tarsus length on the trait scale of mm. P-values were calculated as the proportion of post-warm up draws that were of the opposite value to the mean estimate of the effect of inbreeding

#### FROH

**FuniW** Note: No windows were significant for mass

Table 30: Genetic architecture of inbreeding depression in bill length using FuniW: Table shows the few 2500-SNP windows that cross the bonferonni corrected signifiance threshold.

| Chromosome | Window | estimate | lwr_CI | upr_CI | p_val |
|---|---|---|---|---|---|
| 5B | super_scaffold_18_chunk105 | 1.84 | 0.85 | 2.83 | 1e-14 |

Table 31: Genetic architecture of inbreeding depression in tarsus length using FuniW: Table shows the few 2500-SNP windows that cross the bonferonni corrected signifiance threshold.

| Chromosome | Window | estimate | lwr_CI | upr_CI | p_val |
|---|---|---|---|---|---|
| 1B | super_scaffold_21_chunk90 | -6.50 | -10.29 | -2.82 | 3.333333e-14 |
| 5C | super_scaffold_29_chunk63 | -6.40 | -10.28 | -2.69 | 3.333333e-14 |
| 12 | super_scaffold_3_chunk43 | -6.44 | -10.42 | -2.53 | 3.333333e-14 |
| 19 | super_scaffold_19_chunk17 | 8.13 | 3.75 | 12.48 | 3.333333e-14 |
| 23 | super_scaffold_20_chunk22 | 9.49 | 4.31 | 14.66 | 3.333333e-14 |
